# Integration of Osteoclastogenesis through addition of PBMCs in Human Osteochondral Explants cultured *Ex vivo*

**DOI:** 10.1101/2023.08.08.552563

**Authors:** Esther E.A. Cramer, Bregje W.M. de Wildt, Johannes G.E. Hendriks, Keita Ito, Sandra Hofmann

## Abstract

The preservation of tissue specific cells in their native 3D extracellular matrix in bone explants provides a unique platform to study remodeling. Thus far, studies involving bone explant cultures showed a clear focus on achieving bone formation and neglected osteoclast activity and resorption. To simulate the homeostatic bone environment *ex vivo*, both key elements of bone remodeling need to be represented. This study aimed to assess and include osteoclastogenesis in human osteochondral explants through medium supplementation with RANKL and M-CSF and addition of peripheral blood mononuclear cells (PBMCs), providing osteoclast precursors.

Osteochondral explants were freshly harvested from human femoral heads obtained from hip surgeries and cultured for 20 days in a two-compartment culture system. Osteochondral explants preserved viability and cellular abundance over the culture period, but histology demonstrated that resident osteoclasts were no longer present after 4 days of culture. Quantitative extracellular tartrate resistant acid phosphatase (TRAP) analysis confirmed depletion of osteoclast activity on day 4 even when stimulated with RANKL and M-CSF. Upon addition of PBMCs, a significant upregulation of TRAP activity was measured from day 10 onwards. Evaluation of bone loss trough µCT registration and measurement of extracellular cathepsin K activity revealed indications of enhanced resorption upon addition of PBMCs. Based on the results we suggest that an external source of osteoclast precursors, such as PBMCs, needs to be added in long-term bone explant cultures to maintain osteoclastic activity, and bone remodeling.

**Graphical abstract:** 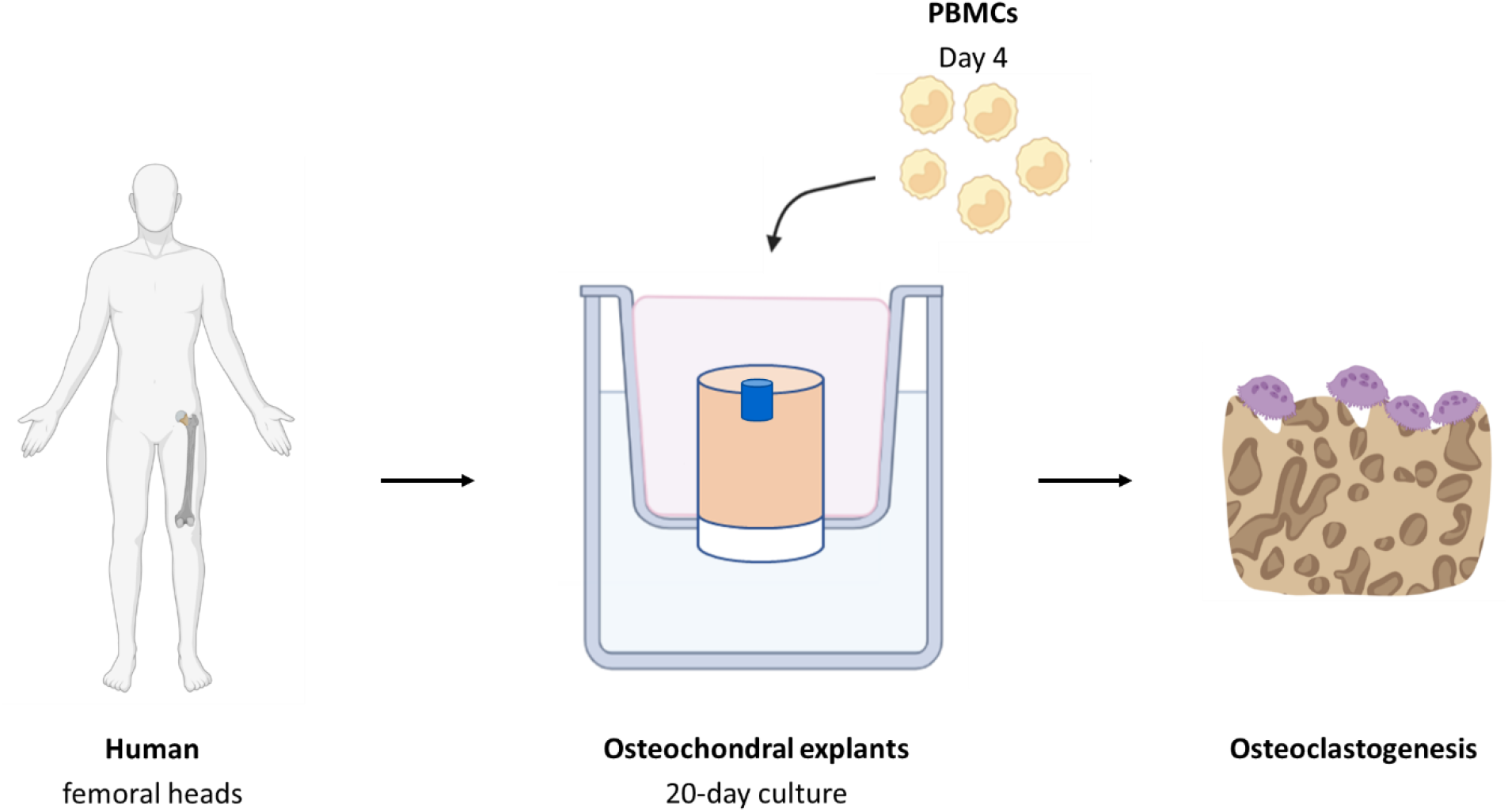

## 1. Introduction

Bone is a hierarchically structured tissue that constantly adapts its architecture to the environment. This continuous process of removal of mineralized bone by osteoclasts followed by the formation of bone matrix by osteoblasts is called bone remodeling. Remodeling processes are regulated through complex interactions between different types of cells, including osteocytes, osteoblasts, osteoclasts, and their progenitors [1]. Bone formation and resorption are often evaluated in preclinical testing of new factors for metabolic bone diseases or novel bone substitutes. Initial screening of novel therapies is regularly performed with simple *in vitro* studies, typically involving 2D single or dual cell cultures to evaluate, for example, communication pathways or toxicity and efficacy [2]. After successful assessment and optimization of new therapies *in vitro*, the preclinical testing process is followed by evaluation in animal models where a poor correlation between *in vitro* test results and *in vivo* outcomes is often observed [3], [4]. A possible explanation for the poor correlation might be the missing interplay between different bone cell types and their spatial arrangement in the original extracellular matrix (ECM) *in vitro*. A 2D culture environment can, for example, result in changes in cell morphology and protein expression because cell polarization is disturbed, thereby leading to different cellular responses compared to *in vivo* [4]–[7]. In addition to the translational issues between *in vitro* and *in vivo* studies, the low success rate, which is about 10%, for novel therapies that make it from beginning of clinical trial to approval for clinical use might partly be attributed to the genetic and physiological differences between the used animal models and humans [8], [9]. To address both translational challenges, human bone explant cultures could give insights in bridging the gap between *in vitro* and *in vivo* experimentation, while also addressing ethical considerations to reduce, refine and replace animal studies (3Rs) [2], [10].

Bone explant cultures could provide a closer representation of the *in vivo* bone environment because cell-cell and cell-matrix interactions are preserved as found *in vivo* [11]. In bone research, explants from porcine, bovine, canine and human origin have been cultured *ex vivo* to assess the effect of biochemical factors and loading on osteocytes as well as to study mechanotransduction pathways and bone formation in static and dynamic culture conditions [12]–[25]. Up until now, studies involving bone explant cultures show a clear focus on improving bone formation and do not incorporate osteoclast activity and resorption [26]. To maintain healthy bone, however, formation is tightly coupled with resorption *in vivo* through pathways acting between osteoclasts and osteoblasts within the basic multicellular units (BMUs) [27]. Specifically, the balance between formation and resorption is regulated by cytokines secreted by osteoblasts, namely receptor activator of NF-κB ligand (RANKL) that stimulates osteoclastogenesis and osteoprotegerin (OPG) that is an inhibitor of RANKL and suppresses osteoclast activity. A few studies evaluated these factors and related pathways in *ex vivo* bone tissue cultures and reported upregulated expression of RANKL after administration of PTH, inflammatory cytokines or bacterial factors [28]–[30]. Moreover, differentiation of osteoclasts was enhanced and increased levels of bone resorption were found upon addition of PTH or bacterial factors S.aureus and LPS to mice explant cultures of flat bones [28], [29], [31]. Another study, using larger bovine trabecular bone core cultures did not show active resorption but stated that osteoclasts were still responsive to drugs retinoic acid and pamidronate [21]. In addition, bovine osteochondral tissue cultures revealed TRAP positive cells after 28 days of culture [32].

For *in vitro* research, it is well-documented that RANKL and macrophage colony stimulating factor (M-CSF) are two essential mediators to initiate differentiation of osteoclast precursors into osteoclasts [33]– [37]. However, these osteoclastic factors are often not incorporated in *ex vivo* cultures since culture medium usually contains supplements to stimulate osteogenic activity, such as dexamethasone, which could even suppress osteoclast activity [21], [38]. Previous research that included osteoclastic factors in a bone explant culture reported that only high concentrations of RANKL (60 ng/ml) and M-CSF (20 ng/ml) did stimulate osteoclastogenesis in mandible slices [28].

Overall, the *ex vivo* studies that assessed osteoclast activity and resorption used animal tissue derived from skull bones, which consist mainly of cortical bone, and do not reflect remodeling levels in long bones of humans containing predominantly trabecular bone [39]. Furthermore, these thin bones have a limited amount of bone marrow, which is the location where hematopoietic stem cells reside, the cells that can differentiate towards osteoclast precursors [40]. Therefore, it would be of interest to study osteoclastogenesis in human trabecular bone explants.

Studying osteoclastogenesis and resorption in long-term explant cultures faces challenges with the average lifespan of mature osteoclasts being 2 weeks and the lack of vascularization *ex vivo*, limiting the source of monocyte-lineage precursors from peripheral blood [41]–[43]. Besides osteoclasts originating from hematopoietic cells in peripheral blood, research showed that also resident precursors in bone marrow are able to differentiate into osteoclasts [44], [45]. Therefore, the aim of this study is to assess the effect of medium supplementation with RANKL and M-CSF on osteoclast activity in human osteochondral explant cultures and to evaluate whether the addition of peripheral blood mononuclear cells (PBMCs), containing monocytes as osteoclast precursors, is needed to enhance osteoclastogenesis. Evaluation of osteoclastogenesis will be performed by a thorough analysis combining qualitative and quantitative techniques, which was often lacking in previous *ex vivo* studies.

## 2. Materials & Methods

### 2.1 Isolation of osteochondral cores

Human femoral heads were obtained from 4 patients undergoing total hip replacements, approved by the Local Medical Ethical Committee (METC, number 16.148). Donor information contained age, gender, and underlying disease. Donor 1: female, 69y, osteoarthritis. Donor 2: female, 62y, osteoarthritis. Donor 3: female, 71y, osteonecrosis. Donor 4: male, 79y, osteoarthritis. After harvesting, femoral heads were wrapped with sterile wet tissues and kept in a sterile container during transportation. All following steps were performed in a sterile environment. Using a hollow-drill (MF Dental, Weiherhammer, Germany) and a custom-made hollow metal tube serving as break off tool, osteochondral cores with a diameter of 10 mm were isolated from the femoral heads (**Figure 1A-B**). The drilling process was performed under constant cooling with sterile PBS (P4417, Sigma-Aldrich) at 4°C containing 2% v/v antibiotic-antimycotic (anti-anti, 15240, Thermo Fisher Scientific). The bone part was cut perpendicular to the longitudinal axis using a bench saw (KS230, Proxxon, Föhren, Germany) to obtain a bone length of approximately 8 mm. It was decided to keep the bone part constant in bone length for each sample. This resulted in osteochondral cores with slightly different final heights because the cartilage layer varied in thickness. A defect (Ø=3 x 2 mm) was drilled, again under constant cooling, in the center of the bone on the trabecular bone site (**Figure 1C-D**) using a custom-made holder. Between steps, explants were stored in sterile PBS with 2% v/v anti-anti. From every donor at least 1 osteochondral core underwent three cycles of freezing at -80°C overnight and thawing at RT to devitalize osteochondral cores to serve as negative control. These controls were cultured in the same way as the other explants. From each donor, between 7 and 14 osteochondral cores were obtained, which were divided among the four different experimental groups. In total, the base medium (BM) and supplemented medium (SM) group included n=10 in each group, the supplemented medium + PBMCs (SM + PBMC) had a sample size of n=11 and devitalized explants (DE) consisted of n=5.

**Figure 1.**
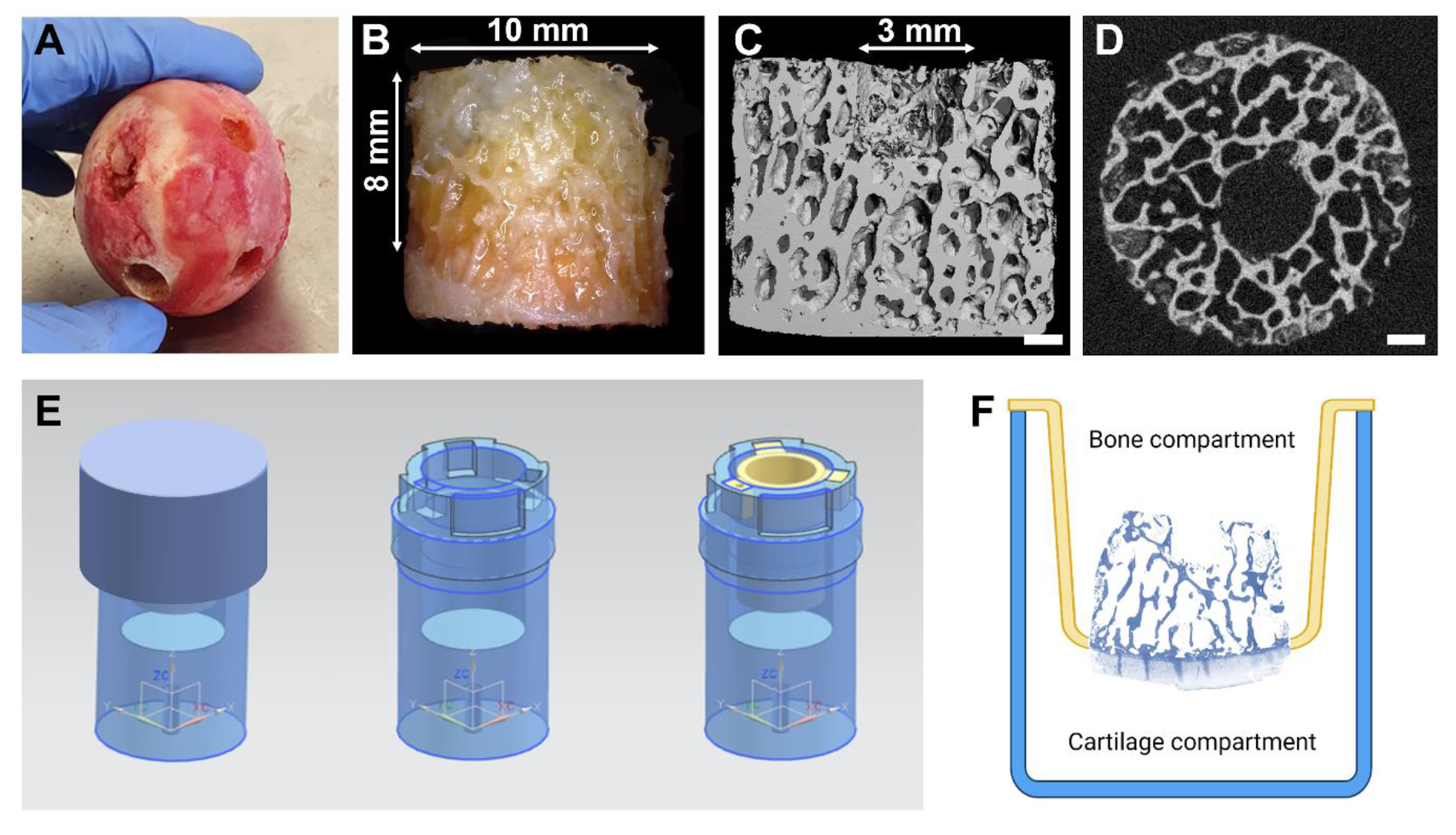
Explant preparation and mounting within the culture chamber system. Human femoral head after removal of osteochondral explants (A). Microscopic image of osteochondral explant with cartilage at the bottom (B). µCT image of side (C) and top (D) view of explant with a defect centered in the explant. Culture chamber system with insert (yellow) to create two compartments (E, F). Scale bar is 1 mm.

### 2.2 Explant culture

After harvesting, each osteochondral core was mounted in an insert using an O-ring (Brammer, Leeds, UK) and placed in a culture chamber creating two separated compartments, with cartilage exposed to medium in the lower compartment and the upper compartment containing the bone part (**Figure 1E-F**). This culture system was closed with a screw thread lid to ensure a sterile environment. This system was adapted from the previous described two-chamber culture system for osteochondral explants in wells-plates [46]. The bottom chamber was filled with 5 ml cartilage medium consisting of α-MEM (41061, Gibco^TM^, Thermo Fisher Scientific), 10% v/v FBS (BCBV7611, Sigma-Aldrich) and 1% v/v anti-anti (15240, Gibco^TM^, Thermo Fisher Scientific). The upper chamber contained 3.5 ml medium which was different for the different experimental groups (**Figure 2**). The base medium group was provided with medium consisting of α-MEM, 10% v/v FBS and 1% v/v anti-anti. The other two conditions, supplemented medium and supplemented medium + PBMCs, received in addition to the base medium 50 ng/ml macrophage colony-stimulating factor (M-CSF, 300-25, PeproTech) and 50 ng/ml receptor activator of nuclear factor κB ligand (RANKL, 310-01, PeproTech). Explants were cultured for 20 days in an incubator at 37°C with 5% CO_2_ and medium was replaced every 2 days in both compartments. With every medium change, supernatant of the bone compartment was saved and stored at -80°C until further analysis.

**Figure 2.**
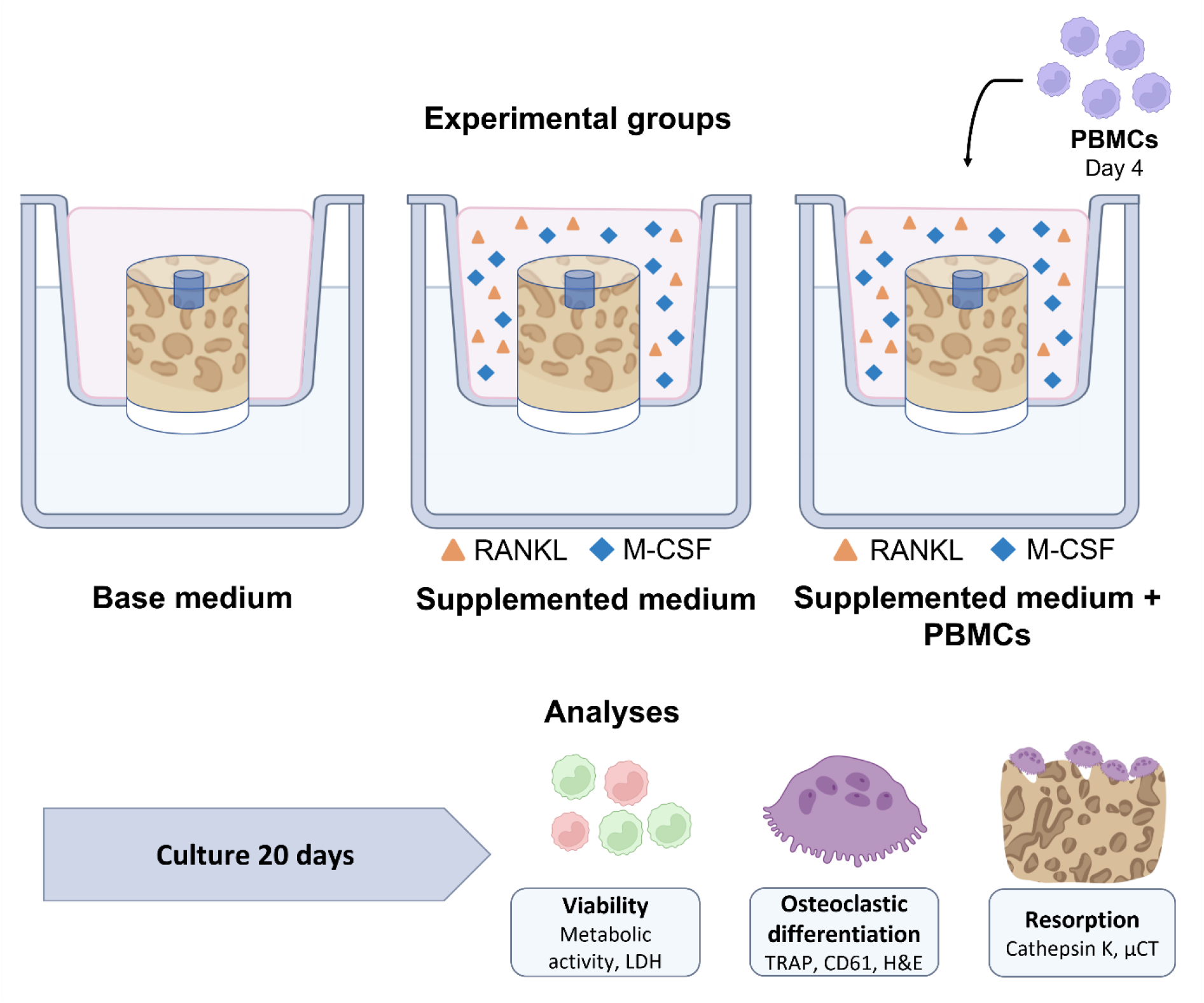
Schematic of experimental set-up with three experimental groups. The bone parts of osteochondral explants received base medium (BM), medium supplemented with RANKL and M-CSF (SM) or medium supplemented with RANKL, M-CSF in combination with PBMCs (SM + PBMC), which were seeded on day 4. After 20 days of culture, multiple types of analyses were performed including viability assessment and evaluation of osteoclastic differentiation and resorption. Abbreviations: peripheral blood mononuclear cells (PBMCs), lactate dehydrogenase (LDH), tartrate resistant acid phosphatase (TRAP), hematoxylin and eosin stain (H&E), micro-computed tomography (µCT). Image created with BioRender.com.

### 2.3 PBMC isolation and seeding

Peripheral blood mononuclear cells (PBMCs) were isolated from human peripheral blood buffy coats of two healthy donors (Sanquin, Eindhoven, The Netherlands, collected under their institutional guidelines and with informed consent per the Declaration of Helsinki). Buffy coats (∼50 ml) were diluted with 0.6% w/v sodium citrate (C7254, Sigma-Aldrich) in PBS (P4417, Sigma-Aldrich) until a final volume of 200 ml. Subsequently, buffy coat-PBS-citrate solution was added in portions of 25 ml to tubes filled with 10 ml Lymphoprep^TM^ (07851, StemCell technologies, Köln, Germany). Tubes were centrifuged (20 min at RT, 800x g, lowest break) to allow for density gradient formation. PBMCs were obtained by collecting the phase formed below the plasma and transferred into PBS-citrate solution. After centrifugation (10 min at RT, 600x g), the cell pellet was resuspended in PBS-citrate solution supplemented with 0.01% w/v bovine serum albumin (BSA, 10735086001, Merck) and centrifuged (5 min at 4°C, 400x g). The resuspension and centrifugation steps were repeated four times to obtain a clear supernatant. PBMCs, containing roughly 10% monocytes, were frozen at 50x10^6^ cells/ml in freezing medium containing 20% v/v FBS (BCBV7611, Sigma-Aldrich) and 10% v/v dimethyl sulfoxide (DMSO, 1.02952.1000, VWR) in RPMI-1640 (RPMI, A10491, Gibco^TM^, Thermo Fisher Scientific), and stored in liquid nitrogen until further use.

### 2.4 Seeding of PBMCs

On day 4 of culture, one experimental group received PBMCs. PBMCs were thawed and resuspended in medium consisting of RPMI, 10% v/v FBS and 1% v/v penicillin-streptomycin (P/S, 15140122, Gibco^TM^, Thermo Fisher Scientific). Cells were centrifuged and resuspended in final culture medium (αMEM, 10% v/v FBS, 1% v/v anti-anti, 50 ng/ml RANKL and 50 ng/ml M-CSF) at a concentration of 100x10^6^ cells/ml. Bone medium was removed from the upper chamber and 250 µl cell solution was added dropwise, in two portions of 125 µl, into the defect of the explants resulting in the addition of 25x10^6^ PBMCs/explant. Cells were allowed to attach for 20 minutes at 37°C before 500 µl culture medium was added. After another hour medium was filled up to 3.5 ml and the culture was continued.

### 2.5 Measurement of cell metabolic activity

To determine cell metabolic activity, AlamarBlue^TM^ (A13262, Thermo Fisher Scientific) was added directly to the culture medium in the bone compartment of the osteochondral explants at a concentration of 10% v/v and incubated for 4.5 hours at 37°C, 5% CO_2_. After incubation, 100 µl of the medium of the upper compartment was pipetted in a black 96-wells assay plate and fluorescence was measured with a plate reader (SynergyTM HTX, Biotek, Winooski, VT, USA) at excitation 530/25 nm, emission 590/35 nm. Measured fluorescent values were corrected for fluorescent values of medium samples of each respective group cultured without explants. After the assay, samples were washed two times with sterile PBS (P4417, Sigma-Aldrich) before fresh medium was added. The AlamarBlue^TM^ assay was performed on day 6 and day 14 of culture on all samples.

### 2.6 Cell death measurement

Lactate dehydrogenase (LDH), an indicator for cell death, was quantified in the culture supernatant taken every 2 days from all samples. Medium samples from day 2 and day 4 had to be diluted 1:1 in PBS (P4417, Sigma-Aldrich) because of the higher levels of cell death in the first days after harvesting caused by the isolation procedure. Medium supernatant, 100 µl, was mixed with an equal amount of LDH reaction mixture, which was prepared according to the manufacturer’s instructions (11644793001, Roche, Sigma-Aldrich). Absorbance was measured directly after mixture and then every 10 minutes to ensure values were within standard curve range using a plate reader (Synergy^TM^ HTX, Biotek, Winooski, VT, USA) at 492 nm. Standards with known NADH concentrations (10107735001, Roche, Sigma-Aldrich), ranging between 0 and 0.75 µmol/ml, were used to calculate the LDH activity per explant.

At the end of culture, on day 20, LDH was determined in both the supernatant to quantify cell death as well as in the bone part of lysed explants as a measure for the remaining living cells. To measure for living cells, osteochondral explants were removed from the insert and cut into one half and two quarters. For both quarter pieces cartilage was removed to evaluate solely the bone part. One quarter was washed two times with PBS (P4417, Sigma-Aldrich) to remove remaining medium, weighted and incubated in 2 mL lysis solution consisting of 1% v/v Triton-X-100 (108603, Sigma-Aldrich) + 5 mM MgCl_2_ (M2393, Sigma-Aldrich) in UPW for 2 hours at RT. Using an ultrasonic disintegrator (Soniprep 150, MSE, France), samples were disintegrated 3 times at 23 KHz with an amplitude of 12 µm for 15 seconds to lyse the remaining cells [47]. After centrifugation at 3000 g for 10 min, LDH assay was performed in a similar way as for supernatant.

### 2.7 DNA quantification

To evaluate the amount of cells that remained in the bone part of explants after 20 days of culture, DNA content was determined. Osteochondral explants were cut as described in section 2.6. A quarter piece with removed cartilage was washed twice with PBS (P4417, Sigma-Aldrich), weighed, and placed into a tube containing 2 mL sterile water. Samples underwent 3 freeze-thaw cycles from -80°C to RT followed by 3 rounds of sonication (Soniprep 150, MSE, France) for 15 sec at 23 KHz with an amplitude of 12 µm and placement on ice between the cycles. After centrifugation at 3000 g for 10 min, DNA content was determined using Qubit^TM^ dsDNA HS Assay Kit (Invitrogen, USA) according to the manufacturer’s instructions. Briefly, 10 µl sample was added to 190 µl of working solution prepared according to the manufacturer’s instructions and measured using the Qubit Fluorometer.

### 2.8 Histological visualization of osteoclasts

Osteoclast-like cells were visualized by staining for tartrate resistant acid phosphatase (TRAP). Directly after isolation (day 0) and after 4 days of culture in medium supplemented with RANKL and M-CSF (without additional PBMCs seeded), osteochondral explants were fixed overnight in 10% neutral buffered formalin at 4°C. Explants were washed in PBS (P4417, Sigma-Aldrich) and immersed in decalcification solution consisting of 10% w/v EDTA (E4884, Sigma-Aldrich) in water at pH 7. This solution was changed 3 times a week for a period of 5 weeks. After decalcification, samples were washed with water to remove remaining EDTA, and subsequently dehydrated through graded ethanol series and xylene in an automatic tissue processor (Microm STP-120, Thermo Scientific) and embedded in paraffin (P3558, Sigma-Aldrich). Sections of 8 µm were sliced using a microtome (Leica RM2255, Germany), attached to Poly-L-Lysine coated microscope slides (Thermo Fisher Scientific) and dried overnight. Slides were deparaffinized in xylene followed by decreasing series of ethanol (3x 100%, 96% and 70%), hydrated and stained for TRAP as follows: Substrate solution was prepared by adding 1 mL Ethylene Glycol Monoethyl Ether (E2632, Sigma-Aldrich) to 20 mg naphthol AS-MX phosphate (N4875, Sigma-Aldrich). Staining solution was created by mixing 0.06% w/v fast red violet LB salt (F3381, Sigma-Aldrich) and 0.5% v/v substrate solution in basic incubation medium, which consisted of 1.14% w/v L- (+) tartaric acid (T6521, Sigma-Aldrich), 0.92% w/v sodium acetate anhydrous (W302406, Sigma-Aldrich) and 0.28% v/v glacial acetic acid (A6283, Sigma-Aldrich) in water. Slides were placed in pre-warmed TRAP staining solution and incubated for 45 min at 37°C. After rinsing in distilled water, samples were counterstained in 0.01% w/v fast green (F7258, Sigma-Aldrich) for 2 min. Slides were air dried, dipped in xylene and mounted with Entellan (107961, Sigma-Aldrich) before imaging under a bright field microscope (Zeiss Axio Observer Z1, Germany).

### 2.9 Visualization of tissue and cellular components in explants

To examine tissue components including cell nuclei and ECM within osteochondral explants, hematoxylin and eosin (H&E) staining was performed. After 20 days of culture, half of the osteochondral explant was fixed, decalcified, dehydrated, embedded in paraffin, and sectioned as described in section 2.8. Slides were deparaffinized in xylene followed by decreasing series of ethanol (3x 100%, 96% and 70%), hydrated and stained with Mayer’s hematoxylin (MHS16, Simga-Aldrich) and Eosin Y (HT110216, Sigma-Aldrich) according to the manufacturer’s instructions. After dehydration in graded series of ethanol (70%, 96%, 3x 100%) and xylene, stained sections were mounted with Entellan (10796,1 Sigma-Aldrich) and imaged using a bright field microscope (Zeiss Axio Observer Z1, Germany). Day 0 samples from each donor were stained with H&E as reference for the start of culture.

### 2.10 Immunofluorescent visualization of osteoclasts

To evaluate the presence of osteoclast-like cells after the 20-day culture period, sections were stained for CD61, also known as integrin-ß3, which is a marker present in osteoclasts. Prepared paraffin sections were deparaffinized and hydrated as described in section 2.8 and stained for CD61and DAPI. Briefly, sections were immersed in 1x citrate buffer (S1699, Dako) overnight at 60°C for antigen retrieval. After a washing step in 0.05% v/v Tween (Tween 20, 822184, Merck) in PBS (P4417, Sigma-Aldrich), samples were blocked in 5% v/v normal goat serum in PBS for 1 hour. Primary antibody solution (orb248939, 1:100, Biorbyt) was incubated overnight at 4°C. Secondary antibody solution (A21240, 1:200, Thermo Fisher Scientific) was incubated for 1 hour at RT followed by DAPI staining, 0.1 µg/ml, for 10 min. Images were acquired with a laser scanning microscope (Leica TCS SP8X).

### 2.11 Measurement of tartrate resistant acid phosphatase activity

To examine differentiation towards osteoclasts, tartrate resistant acid phosphatase (TRAP) activity was quantified in the culture medium that had been saved every 2 days. P-nitrophenyl phosphate buffer was prepared by mixing 1 mg/ml p-nitrophenyl phosphate disodium hexahydrate (71768, Sigma-Aldrich) and 30 µl/ml tartrate solution (3873, Sigma-Aldrich) with stock buffer consisting of 0.1 M sodium acetate (S7670, Sigma-Aldrich) and 0.1% v/v Triton X-100 (108603, Sigma-Aldrich) in PBS (P4417, Sigma-Aldrich). Medium supernatant (10 µl) was added to p-nitrophenyl phosphate buffer (90 µl) and incubated for 90 min at 37°C. To stop the conversion of p-nitrophenyl phosphate into p-nitrophenol, 0.3 M NaOH (100 µl) was added. Absorbance was measured at 405 nm using a plate reader (Synergy^TM^ HTX, Biotek, Winooski, VT, USA). Standards with known p-nitrophenol concentrations ranging between 0 and 0.9 µmol/ml were used to calculate the TRAP activity per explant. Samples of medium cultured without explants were measured for each respective group as reference for background activity in medium.

### 2.12 Measurement of cathepsin K activity

Analysis of resorption by osteoclasts was measured by cathepsin K activity, a protease involved in bone resorption. Cathepsin K was quantified in culture medium which had been collected every 2 days. Substrate working solution was prepared by mixing 1% v/v of 10 mM Z-LR-AMC (BML-P229, Enzo Life Sciences) in DMSO (1.02952.1000, VWR), into 0.1 M sodium acetate (S7670, Sigma-Aldrich) buffer solution containing 4 mM EDTA (E4884, Sigma-Aldrich) and 4 mM dithiothreitol (DTT, RL-1020, Iris Biotech, Germany) at pH 5.5. A volume of 50 µl culture medium was added to 50 µl substrate working solution and incubated for 45 min at 37°C. Fluorescent intensity was measured with a plate reader (SynergyTM HTX, Biotek, Winooski, VT, USA) at excitation 360/40 nm, emission 460/40 nm. Standards with known AMC (A9891, Sigma-Aldrich) concentrations, ranging between 0 and 2 µM, were used to calculate the cathepsin K activity per explant. Samples of medium cultured without explants were measured for each respective group as reference for background activity in medium.

### 2.13 Measurement and visualization of bone gain and loss

Micro-computed tomography (µCT) was used to determine bone gain (formation) and loss (resorption). On day 1, 10 and 20, osteochondral explants, including the devitalized samples, were scanned using micro-computed tomography (µCT100, Scanco Medical, Brüttisellen, Switzerland). Culture chambers were specifically designed to allow µCT scanning during culture without the need to take samples out of their sterile culture environment and to ensure similar orientation of the sample each time. Scanning was performed at an energy level of 55 kVp, intensity of 200 µA and integration time of 200 ms, AL filter of 0.5 mm and a voxel size of 16.4 µm.

For longitudinal image analysis, images of day 20 scans were registered on images of previous scans, either day 10 or day 0, using an automated algorithm in the IPLFE software (Scanco Medical). Voxels appearing only in the first scan were considered as loss and voxels present only in the follow-up scan as gain. Bone gain and loss were determined by finding clusters of ≥ 5 voxels that had a difference in attenuation coefficient between both scans ≥ 225 mgHA/cm^3^. These thresholds were chosen based on literature [48], [49] and based on registration of one sample that was scanned repeatedly on the same day to examine noise levels. With these thresholds applied, noise levels below 1% of the total volume were obtained. Subsequently, the bone was segmented at a threshold of 450 mgHA/cm^3^ and a Gaussian filter with sigma 0.8 and support 1. Bone gain (orange) and loss (purple) were visualized with color-coded images and quantified as percentage of the total bone volume. Percentages of bone loss were subtracted from bone gain to calculate net bone gain, which further reduced noise inclusion.

### 2.14 Statistical analyses

Statistical analyses were performed, and graphs were prepared in GraphPad Prism (version 9.3.0, GraphPad, La Jolla, CA, USA). Data was tested for normal distribution using Shapiro-Wilk tests and homogeneity of variances was assessed from the residual plots. When these assumptions were met, a parametric test was performed. To compare between experimental groups on a specific timepoint a t-test or one-way ANOVA was performed, which was followed by Tukey’s post-hoc analysis. If assumptions of normality and equal variances were not met, a non-parametric Kruskal-Wallis test followed by Dunn’s multiple comparisons test was performed to compare between experimental groups. To compare within a group over time, repeated measures ANOVA was performed, followed by Tukey’s post hoc test. Data is represented as mean ± standard deviation. A p-value of <0.05 was considered statistically significant.

## 3. Results

### 3.1 Osteoclasts did not survive in *ex vivo* culture

Osteochondral explants that were directly fixed after isolation revealed osteoclasts spread throughout the entire bone (**Figure 3A**). Distinct areas of active resorption were observed in trabecular bone (**Figure 3B**) as well as in the subchondral bone (**Figure 3C**) with high numbers of osteoclasts in all donors. After 4 days of culture, osteoclasts were absent as no cells expressing TRAP were visible, not even in samples supplemented with RANKL and M-CSF (**Figure 3D-F**).

**Figure 3.**
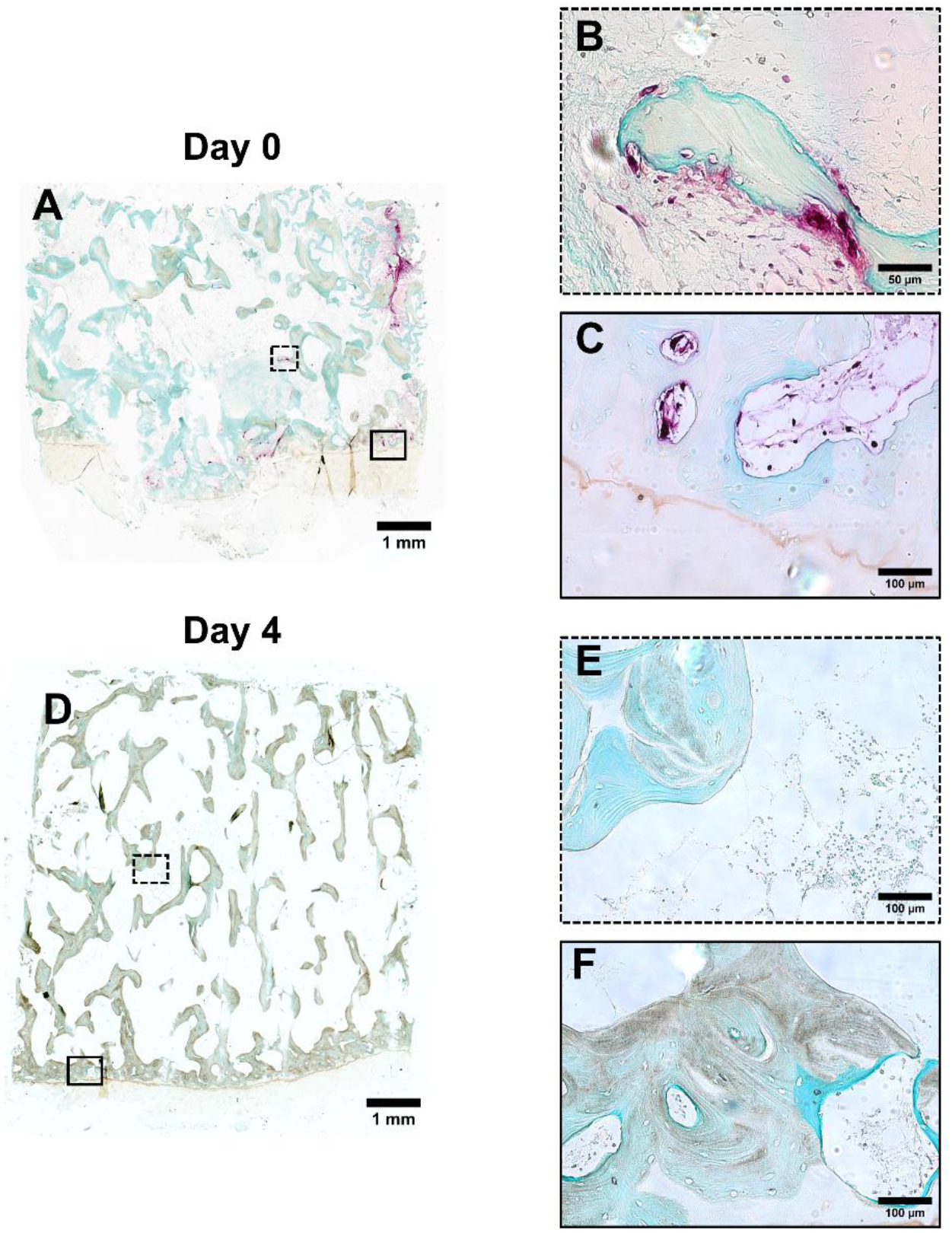
Osteoclasts did not survive within explants cultured *ex vivo*. Representative histological images of osteoclasts (red) in human osteochondral explants stained for TRAP. Osteoclasts were present directly after isolation, day 0 (A-C), and disappeared after 4 days of culture in medium supplemented with RANKL and M-CSF (D-F). Abbreviation: tartrate resistant acid phosphatase (TRAP).

### 3.2 Explants preserved viability during culture

Directly after isolation, high levels of cell death were found for all samples, measured by a high LDH activity in the supernatant that decreased towards day 6 of culture (**Figure 4A**). Stable values of cell death were reached from day 6 onwards with significantly lower cell death levels at the end of culture compared to day 2 and 4 in all groups. Notably, when PBMCs were added to the explants on day 4, mean LDH activity remained constant for this group between day 4, 6 and 8, while the other conditions showed a decrease in cell death in this period (**Figure 4B**). These levels of cell death were, however, not significantly larger upon addition of PBMCs compared to the groups without PBMCs suggesting that most of the added cells attached.

**Figure 4.**
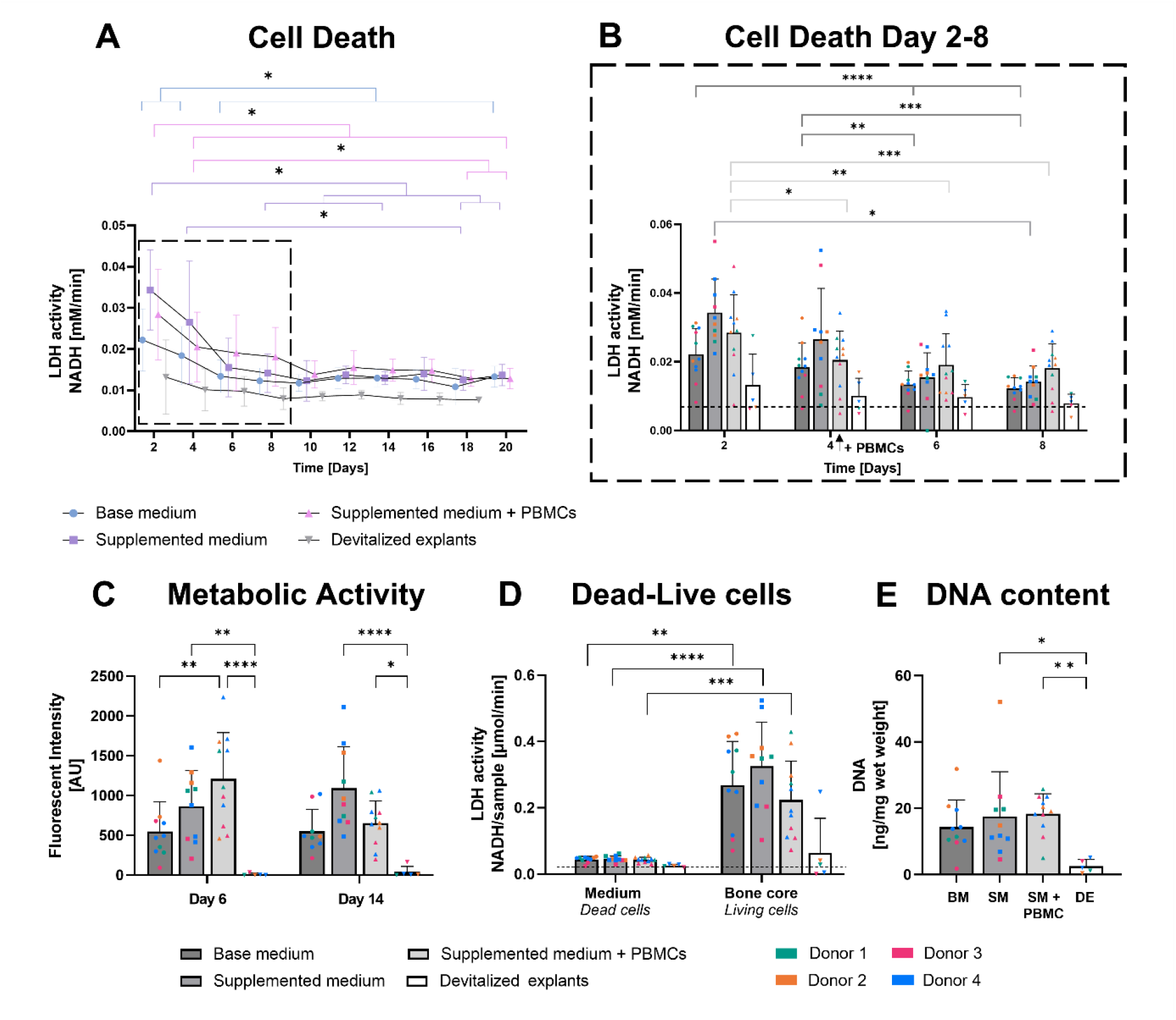
Osteochondral explants remained viable over the culture period. Increased amounts of cell death, measured by LDH release in supernatant, were detected in the first days of culture and declined towards stabilized levels from day 8 onwards (A-B). Zoom of day 2, 4, 6, 8 with the decrease in cell death measured in all groups with living explants (B). Explants maintained metabolic activity on day 6 and 14, whereas devitalized explants were not metabolically active (C). After cell lysis on day 20, explants showed increased levels of LDH activity (living cells) compared to LDH activity in culture medium (dead cells) (D). On day 20, comparable DNA levels were revealed for cultured explants in the three experimental groups being enlarged compared to devitalized explants serving as negative control (E). Data represents mean ± SD (\**p*<0.05, \*\**p*<0.01, \*\*\**p*<0.001, ****p<0.0001). Dashed line indicates LDH activity of medium cultured without explants. Abbreviations: lactate dehydrogenase activity (LDH), peripheral blood mononuclear cells (PBMCs), base medium (BS), supplemented medium (SM), supplemented medium + PBMCs (SM + PBMC), devitalized explant (DE).

At the end of culture, the bone part of the explants was lysed to measure LDH activity derived from vital cells. For all 3 culture conditions, the amount of living cells present at the end of culture was significantly higher, about six times, compared to cell death measured in the supernatant (BM: p=0.002, SM: p<0.0001, SM+PBMC: p=0.0003) (**Figure 4D**). Devitalized explants, used as negative control, showed extra- and intracellular LDH activity in the range of culture medium, which has some LDH activity because of serum supplementation.

Cell metabolic activity was preserved over time for the base medium and supplemented medium group, while it showed a decreasing trend for the group with PBMCs in supplemented medium (**Figure 4C**). On day 6, metabolic activity was significantly higher in the group with PBMCs compared to explants subjected to only base medium (p=0.0026). Since medium was not refreshed before measuring metabolic activity, the increased activity was probably an effect of the addition of 25 million PBMCs on day 4.

Similar levels of DNA were found for the 3 culture conditions at the end of culture (**Figure 4E**), which was in line with LDH and metabolic activity findings demonstrating similar behavior regarding explant viability for all conditions. Devitalized explants revealed little to no metabolic activity and low DNA levels, both significantly lower compared to the conditions with medium supplementation (p=0.032) and PBMCs (0.002).

Visualization of cells and ECM within osteochondral explants at the end of the culture period revealed well-preserved trabecular structures with a comparable presence and distribution of cells for all culture conditions. Cellular structures with intact nuclei were predominantly observed in the subchondral bone and at the edges of the bone surface (**Figure 5A-C**). Osteocytes were preserved in most parts of the trabecular bone (**Figure 5B, C, E**). In central areas of the trabecular bone, regions missing bone surface cells, but containing mainly fat tissue were seen (**Figure 5E**). No signs of increased amounts of cells in explants that received PBMCs were observed visually. However, there were prominent differences between and within donors complicating comparison between explants and culture conditions. A different donor demonstrated high bone marrow cellularity, less fat tissue, and abundance of blood vessels in central regions (**Figure S2A-E**). In addition, variation within a single donor was also present with marrow spaces consisting of only fat tissue and locations that show high bone marrow cellularity. This corresponds to the large spread in datapoints noted from the individual samples in the quantitative outcomes of LDH activity, metabolic activity and DNA content. When each donor was compared to its day 0 situation where remodeling processes with active osteoclasts, osteoblasts and embedding osteoblasts were identified (**Figure S1**), bone marrow cellularity was reduced, and the presence of bone surface cells was decreased and limited (**Figure 5, S2A-E**). Cartilage tissue preserved cells with similar cell morphology and appearance of nuclei compared to uncultured samples (**Figure 5D, S2I**). Histology confirmed absence of living cells in devitalized explants (**Figure S2F-H**).

**Figure 5.**
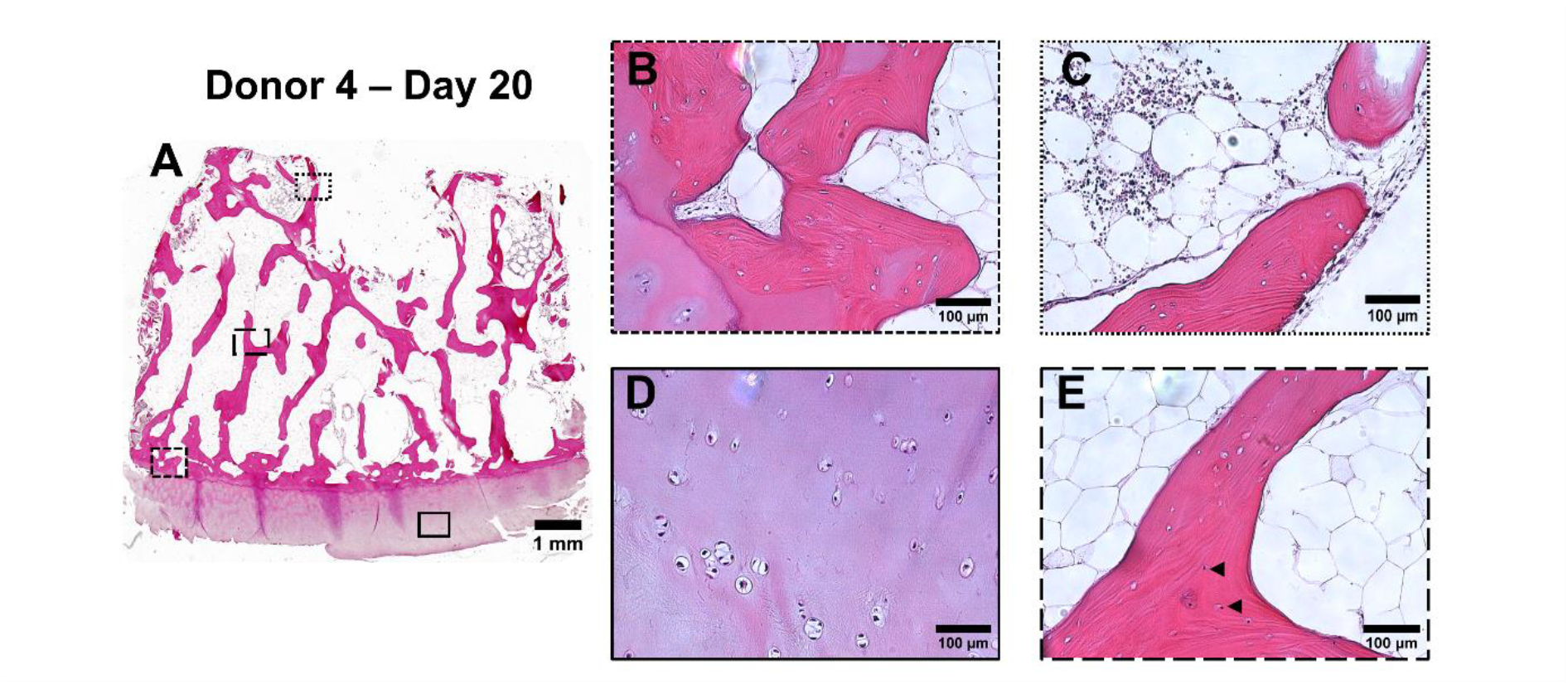
Well-preserved trabecular structures and diverse cell presence and cell structures throughout the explant after culture. Overview of histological images of cross sections of osteochondral explants after 20 days of culture stained with H&E (A). Subchondral bone with osteocytes and marrow cells (B). Location at the edge of the bone part with high marrow cellularity (C). Preserved structural integrity of chondrocytes in cartilage (D). Center of bone part with osteocytes (arrowheads) in lacunae and marrow spaces filled with fat cells (E).

### 3.3 Osteoclastic differentiation was induced in explants upon addition of PBMCs

Adding PBMCs to the explants significantly increased TRAP activity between day 8 and 10 (p=0.019) and TRAP activity continued increasing towards the end of culture for all donors (**Figure 6C**). Compared to the lowest TRAP activity on day 4 (0.62 ± 0.33 µM/min), TRAP activity was three-fold larger on day 20 (1.80 ± 0.61 µM/min) (Day 4-20: p<0.0001), which approached initial values measured on day 2. Explants cultured in all three conditions showed an initial decrease in TRAP activity level between day 2 and day 4 (**Figure 6A-C**), which confirms the histological TRAP activity results where no osteoclasts were seen after 4 days of culture. Interestingly, the only male donor, donor 4, started already with low TRAP activity levels at day 2. As expected, explants cultured in base medium remained at low TRAP activity values for the entire culture (**Figure 6A-B**). Moreover, supplementation with RANKL and M-CSF was not sufficient to stimulate osteoclast differentiation of the naturally present cells. Osteoclast differentiation was only induced upon addition of PBMCs in combination with medium supplementation (**Figure 6B-C**).

**Figure 6.**
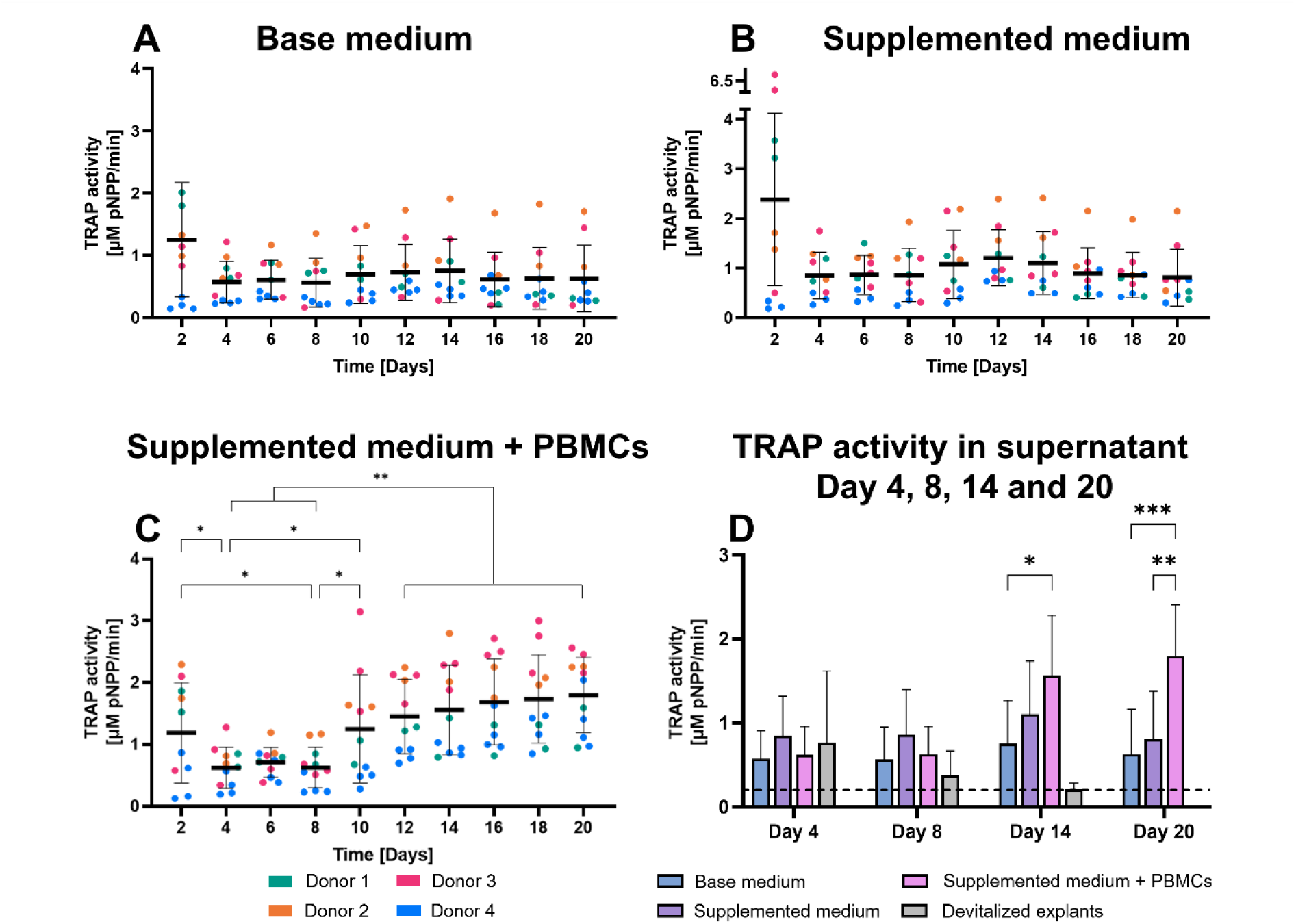
Addition of PBMCs resulted in upregulation of osteoclastic differentiation, measured by TRAP activity in the supernatant, in explants. TRAP levels declined in the first days of culture and remained low for base medium samples (A). Medium supplementation with RANKL and M-CSF did not induce osteoclastic differentiation from resident cells (B). Explants receiving medium supplemented with RANKL and M-CSF in combination with PBMCs, seeded on day 4, demonstrated an increase in TRAP activity from day 10 onwards (C). Comparison between different groups on day 4, 8, 14 and 20 (D). Dashed line is TRAP activity measured in medium cultured without explants. Data represents mean ± SD (*p<0.05, **p<0.01, ***p<0.001). Abbreviations: tartrate resistant acid phosphatase (TRAP), peripheral blood mononuclear cells (PBMCs).

TRAP activity on day 4 and 8 was not significantly different among the three culture conditions of living explants when compared to devitalized explants (**Figure 6D**), indicating the absence of active osteoclasts. Furthermore, the increase in TRAP activity over time seen in the group with PBMCs resulted in significantly higher TRAP levels compared to base medium explants on day 14 and 20 (Day 14: p=0.017, Day 20: p=0.0007), as well as compared to supplemented medium explants on day 20 (p=0.0016).

Only in the explants that received PBMCs, large multinucleated cells close to the bone surface, resembling osteoclast morphology, were observed after 20 days of culture (**Figure 7C**). These histological findings supported the quantitative TRAP activity results. In contrast to day 0 samples, where typical clusters of giant neighboring osteoclasts were found on the bone surface, osteoclast-like cells appeared to be smaller and in less abundant amounts at the end of culture (**Figure S3A-B**). Without PBMCs, cells exhibiting osteoclast characteristics, including multinucleation and being located on the bone surface, were not noticed. A few multinucleated cells, possibly deteriorating as they lack a clear cell membrane, and large cells with multiple lobes, likely megakaryocytes, were occasionally observed (**Figure 7A-B**). When stained for markers specific for osteoclasts, CD61 and TRAP, these markers were only detected in cells within explants that received PBMCs, but not in the explants cultured without PBMCs in either base medium or supplemented medium (**Figure 7D-I, Figure S3C-D**).

**Figure 7.**
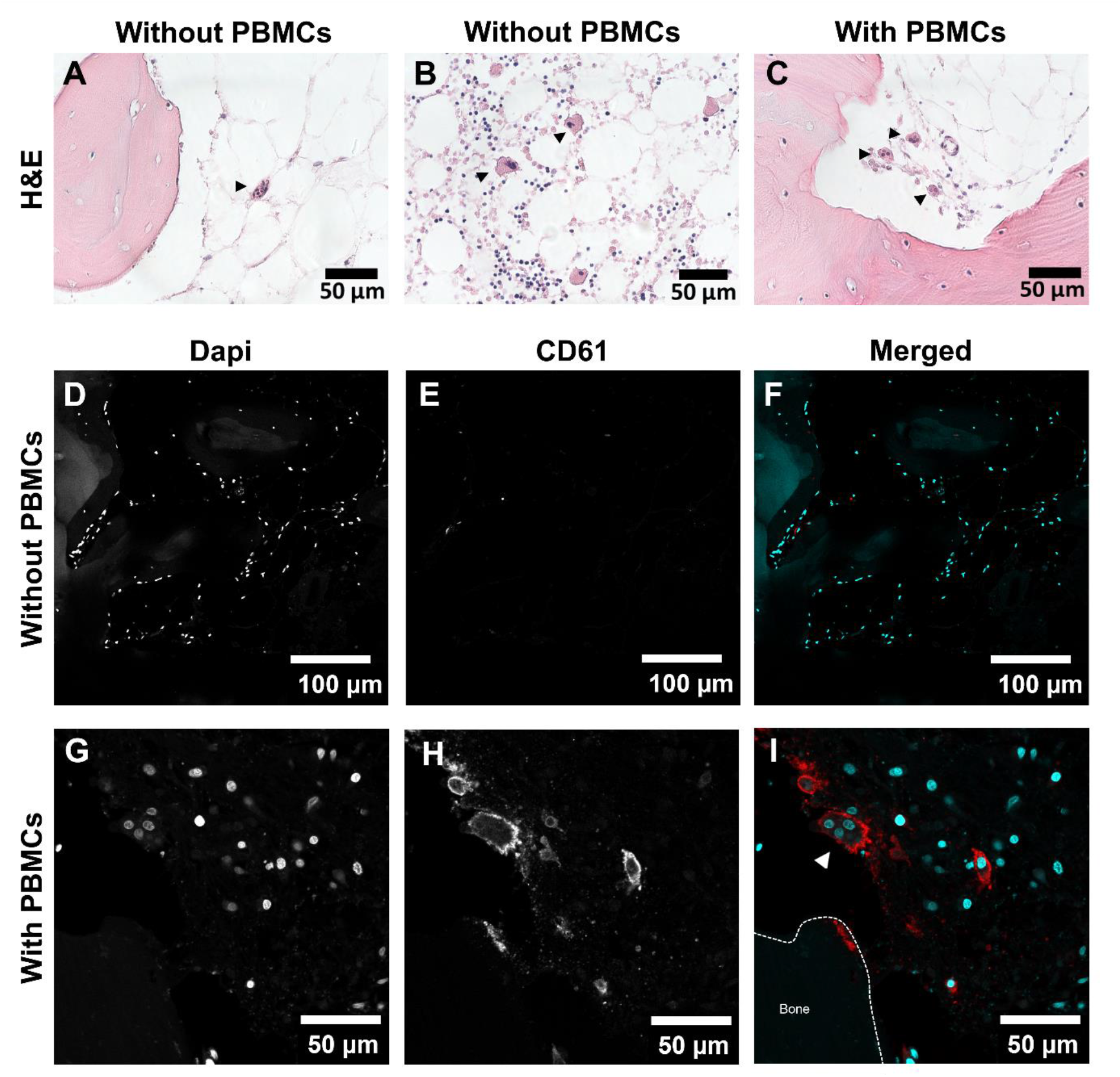
Cells with morphological characteristics specific for osteoclasts, i.e. multiple nuclei, located on the bone surface and positive for a marker specific for osteoclasts, were found only in explants that received PBMCs after 20 days of culture. Histological images of cross sections of osteochondral explants at the end of culture stained with H&E (A-C). In samples without addition of PBMCs, the presence of multinucleated cells without a distinct cell border (A, black arrowhead) and large cells with lobes of nuclei (B, arrowheads) were observed, which can neither be identified as osteoclasts. When PBMCs were added multinucleated cells with a clear cell border close to the bone surface, identified as osteoclasts, were observed in explants (C, black arrowheads). Immunofluorescent images showed absence of osteoclast marker CD61, also known as integrin ß3, (E, red in F) and no multinucleated cells displayed by dapi staining (D and cyan in F) in explants without PBMCs (D-F). Upon addition of PBMCs, osteoclast marker CD61 (H, red in I) was revealed in multinucleated cells (G, cyan in I) in explants (G-I). White arrowhead indicates osteoclast-like cell. Abbreviation: peripheral blood mononuclear cells (PBMCs).

### 3.4 Only explants that received PBMCs demonstrated signs of resorption within explants

In line with TRAP activity results, cathepsin K activity in supernatant decreased between day 2 and day 4 for all groups and remained at this level for explants cultured in base and supplemented medium (**Figure 8A-B**). The addition of PBMCs resulted in an increase in mean cathepsin K activity from day 14 onwards (**Figure 8C**). Mean cathepsin K activity was highest on day 20 (21.56 ± 10.2 nM/min), and was significantly larger compared to day 4, 8 and 12 (lowest was day 8: 11.62 ± 3.0 nM/min) (Day 4-20: p=0.042, Day 8-20: p=0.0005, Day 12-20: p=0.013). Although mean cathepsin K level was increased in the PBMC group, some samples from multiple donors were still at low cathepsin K levels when examining individual samples on day 20.

**Figure 8.**
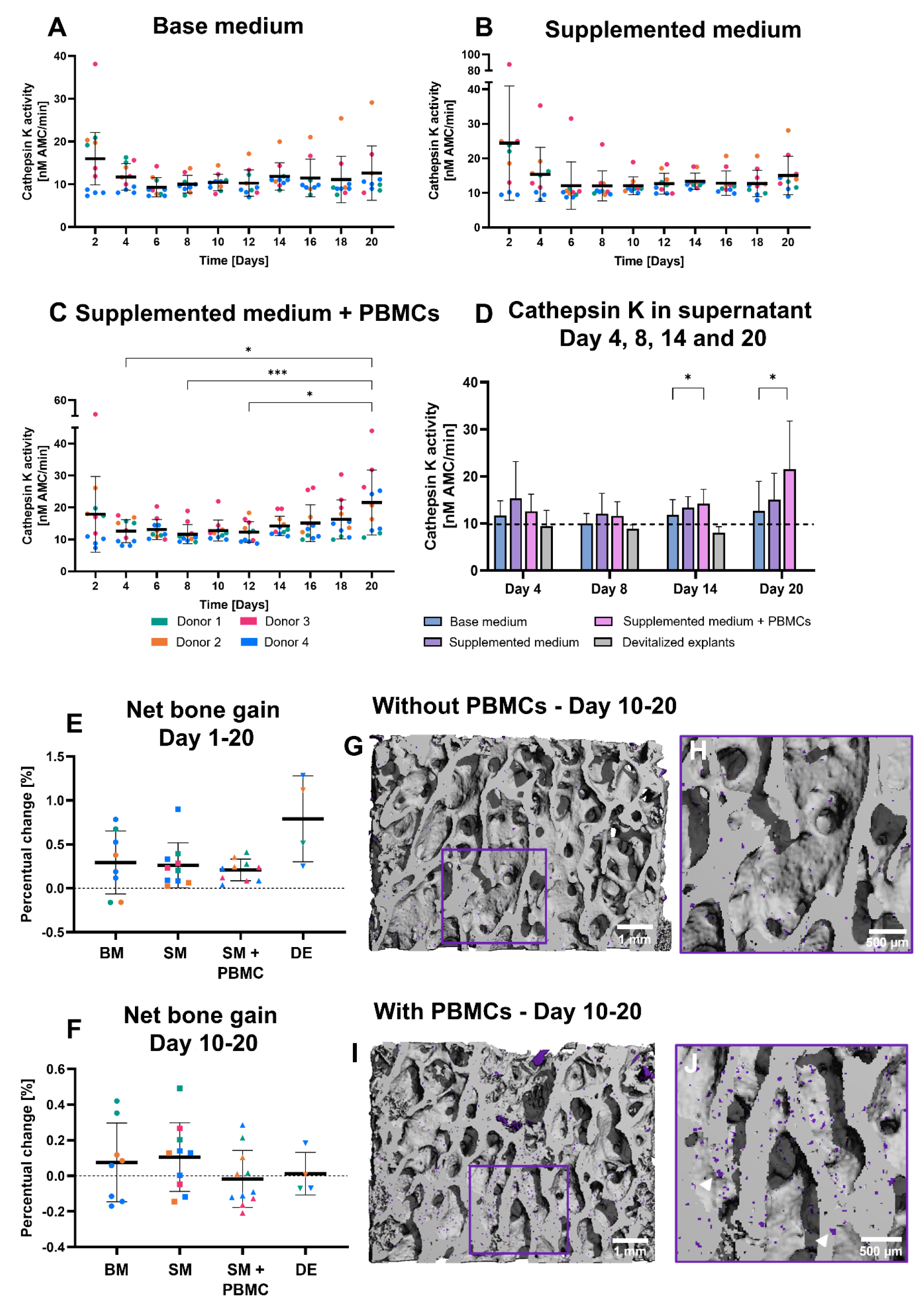
Addition of PBMCs resulted in upregulation of osteoclastic resorption, measured by cathepsin K activity in the supernatant, in explants. Cathepsin K levels declined in the first days of culture and remained low for base medium samples (A). Medium supplementation with RANKL and M-CSF did not induce osteoclastic resorption from resident cells (B). Explants receiving PBMCs, seeded on day 4, and stimulated with medium supplements RANKL and M-CSF demonstrated an increase in cathepsin K activity on day 20 (C). Comparison between different groups on day 4, 8, 14 and 20 (D). Dashed line is cathepsin K activity measured in medium cultured without explants. Net bone gain obtained after µCT registration of day 1 and 20 (E) and day 10 and 20 (F) did not reveal differences between explant groups, although a negative net bone gain was observed between day 10 and 20 for explants receiving PBMCs (F). Visualization of bone loss (purple) of registered images of day 10 and 20 revealed more pronounced spots of bone loss (white arrowhead) on trabeculae when PBMCs were added (H) compared to explants without PBMCs (G). Data represents mean ± SD (*p<0.05, ***p<0.001). Abbreviations: peripheral blood mononuclear cells (PBMCs), base medium (BS), supplemented medium (SM), supplemented medium + PBMCs (SM + PBMC), devitalized explant (DE).

Cathepsin K activity on day 4 and 8 was as low as devitalized explants, indicating the absence of active resorption by osteoclasts (**Figure 8D**). On day 14 and 20, cathepsin K activity was significantly higher in the supernatant of explants that received PBMCs compared to explants cultured without additional PBMCs in base medium (Day 14: p=0.046, Day 20: p=0.013).

Although osteoclast activity was stimulated and therefore increased resorption was expected, analysis of bone remodeling between day 1 and 20 using µCT revealed a small positive mean net bone gain (< 1%) for all explant groups, even in devitalized explants (**Figure 8E, S4E**). This corresponded to the increase in morphometric parameters BV/TV (∼1.8%) and BS/TV (∼1.4%) over the course of the culture found for all explants (**Figure S4C-D**). When evaluating bone remodeling in the second part of culture where monocytes were expected to be differentiated towards osteoclasts, between day 10 and 20, mean net bone gain was lower than in the first period of culture for all groups with no significant differences between the groups (**Figure 8F**). Interestingly, explants with additional PBMCs in supplemented medium revealed even negative mean net bone gain, suggesting a trend towards net bone loss upon addition of PBMCs. Visualization of bone loss showed that, when focusing on areas at the surface of trabeculae, clusters of bone loss were more regularly present in explants that received PBMCs compared to explants lacking PBMCs (**Figure 8G-J**).

## 4. Discussion

Bone explant cultures have the advantage of maintaining tissue specific cells in their native ECM. This provides a unique environment to study bone remodeling processes in which many cell types, such as osteocytes, osteoblasts, osteoclasts, and their progenitors MSCs and monocytes interact in a complex orchestrated way. However, studies involving bone explant cultures show a clear focus on achieving bone formation and have so far neglected looking into osteoclast activity and resorption [26]. Therefore, this study aimed to assess osteoclastogenesis in human osteochondral cores. In addition, it was evaluated if osteoclast differentiation and resorption could be stimulated through the addition of PBMCs, containing monocytes as osteoclast precursor cells.

After 4 days of culture, none of the explants, even upon supplementation with RANKL and M-CSF, showed survival of resident osteoclasts, evaluated by TRAP staining and observed from the decline in TRAP and cathepsin K activity in the supernatant. In contrast, an earlier study demonstrated the presence of osteoclasts, although in low numbers, in thin mandible slices after 14 days of *ex vivo* culture [28]. Within 1 mm mandibular slices TRAP positive cells were identified in the periodontal ligament along the edge of the alveolar bone. With the absence of bone marrow, differences in bone biology between mice and humas, and the small size of the slices not having the issue of avascularity, the mandible explants represent a completely different environment compared to the human bone explants in our study [50]. These factors might explain the differences in osteoclast presence. A study by de Vries-van Melle et al. investigated osteoclast presence in larger osteochondral cores cultured for 4 weeks [32]. They reported preservation of osteoclasts, but histological images showed diminished TRAP staining signals towards the end of culture and lacked contours of large multinucleated cells, questioning the presence of active osteoclasts.

Based on the elevated levels of cell death in the first days observed in the present study, it is suggested that osteoclasts have died. This could be a result of the change from *in vivo* environment to an artificial *ex vivo* environment not having a vascular system and lacking innervation and mechanical loading which could evoke changes in signaling pathways potentially causing cell death. Generally, reduced amounts of loading or unloading result in bone adaptation by increased osteoclastic resorption *in vivo* [51]–[53]. A previous *ex vivo* study showed that explants cultured in near-weightlessness demonstrated reduced levels of bone volume and decreased mechanical properties suggesting an increased resorption, although thorough analysis of osteoclast activity was not performed [54]. On the other hand, *Valentin et al.* cultured bone explants in a perfusion compression bioreactor and described that osteoclasts were still responsive upon drug treatment [21]. Although histological verification of osteoclastogenesis was missing in their study, the incorporation of perfusion to improve nutrient supply, which is missing in our static culture system used, might be a factor necessary to maintain native osteoclastic activity *ex vivo*.

Even when supplementing explants with RANKL and M-CSF, osteoclastogenesis from resident cells could not be detected. This questions the presence and/or reactivity of osteoclast precursors in bone marrow in these explants. As the bone tissue is obtained from donors aged above 60y, the bone is likely to contain yellow marrow, rich in adipocytes and with reduced hematopoietic activity, but the marrow content was not specifically characterized in the explants [55]–[57]. On the other hand, previous research demonstrated that bone marrow cells isolated from human femoral heads of osteoarthritic patients were able to differentiate into viable and functional osteoclasts *in vitro* [58]. It was found that only optimal culture conditions, 25 ng/ml M-CSF for 3 days followed by 50 ng/ml RANKL and 25 ng/ml M-CSF for 11 days, generated large amounts of osteoclasts [58]. In addition, *ex vivo* culture of murine mandibles showed that low concentrations of RANKL (30 ng/ml) and M-CSF (10 ng/ml) did not affect osteoclast numbers, while higher concentrations of RANKL (60 ng/ml) and M-CSF (20 ng/ml) did stimulate osteoclastogenesis from resident cells. The enhanced osteoclastogenesis was even more pronounced after stimulation with inflammatory factors, bacterial LPS and TNFα [28]. In our study, RANKL (50 ng/ml) and M-CSF (50 ng/ml) were concurrently administered from the start of culture and in concentrations different from other literature which might explain the described results. Since these studies highlight the importance of optimized culture conditions to induce osteoclastogenesis from bone marrow cells, it would be interesting to study the effect of different administration patterns and concentrations of RANKL and M-CSF as well as to test different factors such as PTH, vitamin D3, bacterial LPS, TGF-β and TNFα [59]–[62]. Although inflammation-mediated osteoclast activity is not desired in homeostatic bone remodeling it could give insights in the reactivity of resident bone marrow osteoclast precursor cells in the osteochondral explant system.

The combination of medium supplementation with RANKL and M-CSF and addition of PBMCs as a fresh source of monocytes on day 4 of culture induced osteoclastic differentiation within the explants. With supernatant samples taken every two days, osteoclastic differentiation was closely monitored by means of extracellular TRAP activity, an enzyme synthesized and secreted by osteoclasts. Here, increased extracellular TRAP activity was first observed on day 10 and continued towards the end of the culture, which corresponds to the timing of the differentiation process from peripheral blood derived monocytes into osteoclasts as seen *in vitro*, which requires 4-5 days before giant multinucleated cells appear [63]–[66]. In addition, TRAP release has been reported to continue at least up to 16 days [63]. Comparison of TRAP measurements with those of *in vitro* studies with similar amounts of monocytes and comparable culture conditions demonstrates a corresponding range of extracellular TRAP, between 1 and 1.5 µM/min, after 14 and 21 days of culture [64], [67]. At the end of culture, osteoclastic differentiation was confirmed by histology and immunohistochemistry showing multinucleated cells positive for osteoclast markers CD61 and TRAP close to the bone surface. Although levels of TRAP activity in the supernatant reached a similar range at the end of culture as found on day 2, visual differences were observed compared to uncultured explants as osteoclast-like cells seemed smaller with less nuclei and not present in clusters forming borders of osteoclasts occupying substantial parts of the bone surface. A possible explanation for this might be the younger age of PBMC donors compared to the age of the femoral head donors, as nucleation number per osteoclast was found to increase with age [68].

Within the osteochondral explants, which lack vascularization and being depleted of infiltrating blood monocytes, resident precursor cells were not capable of differentiating into osteoclasts, while peripheral blood derived precursors did differentiate into osteoclasts. Several studies reported differences in osteoclastogenic potential and resorption patterns between bone marrow derived and peripheral blood derived osteoclasts and suggested that peripheral blood monocytes have increased osteoclast differentiation potential [59], [69], [70]. There is increasing evidence that osteoclast precursors are released from the bone marrow into the bloodstream, circulate as cell cycle-arrested quiescent osteoclast precursors (QOPs), a population of circulating blood monocytes, and migrate from the capillaries into bone tissues to complete osteoclastogenesis [42], [59], [71], [72]. The concept of circulating QOP as the main source of osteoclast precursors seems in line with the results found in this study that precursors within the added PBMC population were able to differentiate towards osteoclasts. Whether bone marrow osteoclast precursors are present in the marrow and can migrate locally and directly to remodeling bone surfaces remains unclear [42], [73].

Towards the end of the culture the first signs of upregulated osteoclastic resorption, measured by cathepsin K activity in the supernatant, were detected in explants that received PBMCs. Active osteoclasts resorb bone by first releasing acid phosphatases into the resorption lacunae to degrade inorganic matrix and subsequently proteolytic enzymes, such as cathepsin K, to cleave and degrade organic bone matrix [74]. Research has shown that extracellular cathepsin K activity is correlated with extracellular TRAP activity and the resorbed area on dentin [75]. Where cathepsin K measurement is used to gain insight into total resorptive turnover of the explant, histology is generally performed for local visualization of resorption. However, it is challenging to show new resorption *ex vivo* as resorption pits exist already upon isolation of the explants. µCT image registration allows longitudinal quantification and visualization of bone loss in which each sample served as its own reference. Since osteoclast-like cells did not appear as a multitude of cells populating the bone surface but rather as individual osteoclasts spread throughout the explant, large resorption surfaces were not expected. This challenged the use and interpretation of the µCT outcomes since the voxel size is 16.4 µm and only a cluster of 5 voxels, approximately the size of a single resorption pit, was included as gain or loss to limit noise. To further exclude noise and artifacts caused by movement of bone debris that appeared in both gain and loss net bone gain was calculated instead of evaluating solely bone loss (**Figure S4A-B**). Visualization of bone remodeling within the devitalized explants revealed that complete removal of noise could not be achieved as spots of both gain and loss were distributed throughout the bone. However, these spots were mainly localized inside trabeculae while remodeling is expected at the surface (**Figure S4F-G**). Therefore, focus was given to larger clusters of bone loss at the surface. Using this method to interpret µCT outcomes, there was a trend observed towards a slight upregulation of bone resorption in explants that received PBMCs compared to explants lacking PBMCs, which is in accordance with the measurement of cathepsin K activity in the supernatant. It seems that the osteoclasts’ resorptive activities had just started at the end of the culture as not all explants showed signs of increased resorption on day 20. It remains to be elucidated if an increased culture time could achieve higher levels of resorption in all explants.

Quantitative and qualitative outcomes of this study showed that we were able to preserve viability of bone explants over the course of 20 days. Nonetheless, the elevated levels of cell death, measured by extracellular LDH, in the first days of culture indicated a decrease in cell viability. This finding seems inherent to the explantation of bone tissues as cells are exposed to hypoxia and mechanical stress during harvesting procedures [32], [47], [76]. Thus far, it is not reported whether specific types of resident cells die upon explant isolation and which cell types are preserved. Regarding bone cells, this study showed active osteoclasts were absent, while histologically intact osteocytes were present in their lacunae at the end of culture. These osteocytes appeared morphologically comparable to uncultured samples, therefore assuming viable osteocytes. This is in line with Zankovic et al. who found viable osteocytes after 28 days of bone explant culture [77]. It would be of interest to study interactions between the bone cell types remaining within the explants as for example, previous research suggested that living osteocytes inhibit bone resorption *ex vivo* [78]. In their study they found that seeded osteoclasts were active in devitalized calvarial slices whereas in explants with living osteocytes, osteoclasts were disturbed in their actin ring formation [78]. Next to osteogenic cells, bone houses a variety of other cell types, including fibroblasts, pericytes, fat cells and immune cells, which could impact bone remodeling [79], [80]. A limitation of this study was that the femoral heads were obtained from diseased old patients (> 60 y). They might not represent bone remodeling and cell behavior in younger and/or healthy persons. Upon aging, alterations in bone structure as well as in the bone marrow occur, such as decline in bone marrow cellularity and increase in adipocyte content, which in combination with the underlying pathologies could explain high variability in patient tissue as was seen for the donors included here [81]–[83]. Whereas the main underlying disease of the patients was osteoarthritis, one osteonecrotic femoral head was included. From this femoral head, samples were not obtained from the necrotic region as bone tissue is dying and osteoclast activity could differ from healthy regions in an osteonecrotic femoral head [84]. The obtained explants from the osteonecrotic femoral head did not show differences in viability or osteoclast activity compared to explants from other donors. An interesting finding in variability regarding osteoclast activity was that the one male donor did not have initial TRAP activity, while the other female donors did show distinct levels of TRAP in the supernatant at the start of culture. This might be related to increased levels of resorptive activity appearing after the menopause as a result of the decrease in estrogen levels [85], [86], although care should be taken with this suggestion as only one male donor was included and other donor specific aspects, which are unknown, could have had an influence. Donor variability was not only observed between donors but also within a donor comprising locations in the femoral head with high marrow cellularity and areas with low cellularity. This was also reflected in the bone structure with parts containing either extensive and thick or open and thin trabecular structures, which could be explained by structural adaptation upon loading patterns observed within the femoral head causing active and quiescent areas [84], [87], [88]. Furthermore, donors for bone explant tissue were not matched with PBMC donors with respect to sex and age. However, it was demonstrated that PBMCs derived from different donors were capable to differentiate into osteoclast-like cells in all 4 explant donors. Ideally, it is desirable to retrieve blood from the same donor as the explant tissue to have a closer representation of remodeling processes in the bone environment of one single individual. Despite the differences between and within donors, the pattern during the first days was identical in all explants, in all conditions, generating a comparable situation on day 4 i.e., a viable osteochondral core without resident osteoclasts.

The culture system used in this study was comprised of two separate media chambers to allow culturing of bone and cartilage, each in their own compartment. Earlier research using this culture platform to study cartilage defect treatments and cartilage preservation of human osteoarthritic tissue revealed that the bone part remained viable during culture [46], [89]. Here, human tissue from primarily osteoarthritic patients was used, of which it is known that the subchondral bone can be affected and is being highly remodeled [90]. This was confirmed by the abundancy of osteoclasts being present in the subchondral bone directly after isolation. Therefore, it was decided to keep the cartilage-bone interface intact and make use of the two-compartment culture system. Moreover, by preserving the cartilage layer no additional damage from drilling and sawing procedures was generated. Although the two-compartment system complicated the culture set-up and medium changes required additional time compared to culture in a simple well-plate, it allowed each tissue to be exposed to its specific medium and it kept the explants in a fixed orientation during culture, which is important for repeated µCT scanning and subsequent registration.

Although osteoblast activity and bone formation were not specifically assessed in our study, it was demonstrated that devitalized explants acquired mineralized matrix during the culture period. This was probably an effect of precipitation of calcium and phosphate from the medium because no viable cells were detected in these explants (**Figure S4C-E**). This type of volume gain, not caused by cellular activity, should be considered during evaluation as it might lead to over-interpreting bone formation and highlights the importance of including negative controls. Nonetheless, earlier research has shown that osteoblasts survive in bone explants and are able to deposit new matrix, especially under mechanical stimulation [21]. With the incorporation of osteoclastogenesis in those culture systems by adding PBMCs and medium factors RANKL and M-CSF as described here, the tightly coupled processes between osteoblasts, osteocytes and osteoclasts can be studied. Therefore, culture medium should be further supplemented with growth factors necessary to stimulate osteoblast activity such as ascorbic acid, ß-glycerophosphate and optionally dexamethasone. It is important to optimize the combination of growth factors and their concentrations in culture medium to prevent negative effects on either osteoclasts or osteoblasts. For example, there are conflicting results of the effect of dexamethasone on osteoclast differentiation and activity, demonstrating both stimulation of proliferation as well as inhibition of osteoclast formation, which is probably dependent on the provided concentration [38], [86], [87]. Ideally, we get towards a self-sustaining system where resident cells influence each other and achieve a homeostatic state of bone remodeling without the addition of such powerful growth factors [67]. It is expected that in a self-regulating *ex vivo* system it would be easier to detect bone remodeling effects of new therapeutics since cells are not affected in their behavior by the cocktail of growth factors added. In that case, human explant cultures could potentially contribute to preclinical development of bone remodeling drugs, especially when tissue from the elder population is included as this is an important target group with the high occurrence of bone diseases.

## 5. Conclusion

In conclusion, the addition of PBMCs was demonstrated as a successful method to integrate osteoclastic activity, an essential component of bone remodeling, in the *ex vivo* culture of human osteochondral explants. Our study revealed that the static culture condition, even upon supplementation with RANKL and M-CSF, was unable to preserve active resident osteoclasts and did not induce osteoclastogenesis from resident precursor cells. Upon addition of PBMCs, osteoclastic differentiation and initial signs of resorption were demonstrated, suggesting that an additional source of monocytes, such as PBMCs, should be considered when studying osteoclast activity, and bone remodeling, during long-term human *ex vivo* culture studies. Taken together, a first step was taken to incorporate osteoclast activity and resorption in *ex vivo* bone explant cultures. These explant cultures could potentially provide an *ex vivo* model to evaluate novel bone remodeling therapeutics while they could also contribute to better translation between *in vitro* and *in vivo* studies.

## CRediT authorship contribution statement

E.C., K.I. and S.H. contributed to conception and design of the study. E.C., B.d.W. and S.H. contributed to the methodology. E.C. performed experiments, analyzed results, and wrote the draft of the manuscript. J.G.E.H. contributed to tissue acquisition and harvesting. S.H. and K.I. contributed to the supervision. SH acquired funding for this research. All authors contributed to manuscript revision and approved the submitted version.

## Acknowledgement

The authors would like to thank Jurgen Bulsink for manufacturing and maintenance of the culture chambers and tools for tissue isolation. We also gratefully acknowledge the Gravitation Program “Materials Driven Regeneration” (024.003.013), funded by the Ministry of Education, Culture and Science.

## References

[1] A. G. Robling, A. B. Castillo, and C. H. Turner, “Biomechanical and molecular regulation of bone remodeling,” Annu Rev Biomed Eng, vol. 8, no. 1, pp. 455–498, Aug. 2006, doi: 10.1146/annurev.bioeng.8.061505.095721.

[2] S. K. Doke and S. C. Dhawale, “Alternatives to animal testing: A review,” Saudi Pharmaceutical Journal, vol. 23, no. 3. Elsevier, pp. 223–229, Jul. 01, 2015. doi: 10.1016/j.jsps.2013.11.002.

[3] G. Hulsart-Billström et al., “A surprisingly poor correlation between in vitro and in vivo testing of biomaterials for bone regeneration: results of a multicentre analysis,” Eur Cell Mater, vol. 31, pp. 312–322, May 2016, doi: 10.22203/eCM.v031a20.

[4] M. Peroglio, D. Gaspar, D. I. Zeugolis, and M. Alini, “Relevance of bioreactors and whole tissue cultures for the translation of new therapies to humans,” Journal of Orthopaedic Research, vol. 36, no. 1, pp. 10–21, Jan. 2018, doi: 10.1002/jor.23655.

[5] A. Sieberath, E. Della Bella, A. M. Ferreira, P. Gentile, D. Eglin, and K. Dalgarno, “A comparison of osteoblast and osteoclast in vitro co-culture models and their translation for preclinical drug testing applications,” International Journal of Molecular Sciences, vol. 21, no. 3. MDPI AG, Feb. 01, 2020. doi: 10.3390/ijms21030912.

[6] D. W. Hutmacher, D. Loessner, S. Rizzi, D. L. Kaplan, D. J. Mooney, and J. A. Clements, “Can tissue engineering concepts advance tumor biology research?,” Trends Biotechnol, vol. 28, no. 3, pp. 125–133, Mar. 2010, doi: 10.1016/j.tibtech.2009.12.001.

[7] G. D. Prestwich, “Simplifying the extracellular matrix for 3-D cell culture and tissue engineering: A pragmatic approach,” J Cell Biochem, vol. 101, no. 6, pp. 1370–1383, Aug. 2007, doi: 10.1002/JCB.21386.

[8] F. Barré-Sinoussi and X. Montagutelli, “Animal models are essential to biological research: issues and perspectives,” Future Sci OA, vol. 1, no. 4, Nov. 2015, doi: 10.4155/FSO.15.63.

[9] D. Thomas et al., “Clinical Development Success Rates and Contributing Factors 2011-2020,” 2021.

[10] W. Russell and R. Burch, The principle of Humane Experimental Technique. Londen: Methuen, 1959.

[11] S. Marino, K. A. Staines, G. Brown, R. A. Howard-Jones, and M. Adamczyk, “Models of ex vivo explant cultures: applications in bone research.,” Bonekey Rep, vol. 5, p. 818, 2016, doi: 10.1038/bonekey.2016.49.

[12] M. E. Chan et al., “A Trabecular Bone Explant Model of Osteocyte–Osteoblast Co-Culture for Bone Mechanobiology,” Cell Mol Bioeng, vol. 2, no. 3, p. 405, Sep. 2009, doi: 10.1007/S12195-009-0075-5.

[13] E. Takai, R. L. Mauck, C. T. Hung, and X. E. Guo, “Osteocyte Viability and Regulation of Osteoblast Function in a 3D Trabecular Bone Explant Under Dynamic Hydrostatic Pressure,” Journal of Bone and Mineral Research, vol. 19, no. 9, pp. 1403–1410, Jun. 2004, doi: 10.1359/JBMR.040516.

[14] J. Gao et al., “Glucocorticoid impairs cell-cell communication by autophagymediated degradation of connexin 43 in osteocytes,” Oncotarget, vol. 7, no. 19, pp. 26966–26978, May 2016, doi: 10.18632/oncotarget.9034.

[15] F. Li, J. D. Cain, J. Tombran-Tink, and C. Niyibizi, “Pigment epithelium-derived factor (PEDF) reduced expression and synthesis of SOST/sclerostin in bone explant cultures: implication of PEDF-osteocyte gene regulation in vivo,” J Bone Miner Metab, vol. 37, no. 5, pp. 773–779, Sep. 2019, doi: 10.1007/s00774-018-0982-4.

[16] E. H. Davidson et al., “Flow perfusion maintains ex vivo bone viability: a novel model for bone biology,” J Tissue Eng Regen Med, vol. 6, no. 10, pp. 769–776, Nov. 2012, doi: 10.1002/term.478.

[17] E. Lozupone, C. Palumbo, A. Favia, M. Ferretti, S. Palazzini, and F. P. Cantatore, “Intermittent Compressive Osteogenesis and Improves in Bones Cultured Load Stimulates Osteocyte Viability ‘In Vitro,’” 1996.

[18] J. Vivanco, S. Garcia, H. L. Ploeg, G. Alvarez, D. Cullen, and E. L. Smith, “Apparent elastic modulus of ex vivo trabecular bovine bone increases with dynamic loading,” Proc Inst Mech Eng H, vol. 227, no. 8, pp. 904–912, Aug. 2013, doi: 10.1177/0954411913486855.

[19] M. Kogawa et al., “Recombinant sclerostin antagonizes effects of ex vivo mechanical loading in trabecular bone and increases osteocyte lacunar size,” American Journal of Physiology-Cell Physiology, vol. 314, no. 1, pp. C53–C61, Jan. 2018, doi: 10.1152/ajpcell.00175.2017.

[20] K. J. Curtis, T. R. Coughlin, D. E. Mason, J. D. Boerckel, and G. L. Niebur, “Bone marrow mechanotransduction in porcine explants alters kinase activation and enhances trabecular bone formation in the absence of osteocyte signaling,” Bone, vol. 107, pp. 78–87, Feb. 2018, doi: 10.1016/j.bone.2017.11.007.

[21] V. David et al., “Ex Vivo Bone Formation in Bovine Trabecular Bone Cultured in a Dynamic 3D Bioreactor Is Enhanced by Compressive Mechanical Strain,” Tissue Eng Part A, vol. 14, no. 1, pp. 117–126, 2008, doi: 10.1089/ten.a.2007.0051.

[22] V. Mann, C. Huber, G. Kogianni, D. Jones, and B. Noble, “The influence of mechanical stimulation on osteocyte apoptosis and bone viability in human trabecular bone,” Journal of Musculoskeletal Neuronal Interactions, vol. 6, no. 4, pp. 408–417, Oct. 2006.

[23] F. Rupin et al., “Adaptive remodeling of trabecular bone core cultured in 3-D bioreactor providing cyclic loading: An acoustic microscopy study,” Ultrasound Med Biol, vol. 36, no. 6, pp. 999–1007, 2010, doi: 10.1016/j.ultrasmedbio.2010.03.004.

[24] R. Dua, H. Jones, and P. C. Noble, “Designing and validation of an automated ex-vivo bioreactor system for long term culture of bone,” Bone Rep, vol. 14, p. 101074, Jun. 2021, doi: 10.1016/J.BONR.2021.101074.

[25] S. Endres, M. Kratz, S. Wunsch, and D. B. Jones, “Zetos: A culture loading system for trabecular bone. Investigation of different loading signal intensities on bovine bone cylinders,” Journal of Musculoskeletal Neuronal Interactions, vol. 9, no. 3, pp. 173–183, 2009.

[26] E. E. A. Cramer, K. Ito, and S. Hofmann, “Ex vivo Bone Models and Their Potential in Preclinical Evaluation,” Curr Osteoporos Rep, vol. 19, no. 1, p. 75, Feb. 2021, doi: 10.1007/S11914-020-00649-5.

[27] R. Owen and G. C. Reilly, “In vitro Models of Bone Remodelling and Associated Disorders,” Front Bioeng Biotechnol, vol. 6, no. OCT, p. 134, Oct. 2018, doi: 10.3389/FBIOE.2018.00134.

[28] A. J. Sloan et al., “A novel ex vivo culture model for inflammatory bone destruction.,” J Dent Res, vol. 92, no. 8, pp. 728–34, Aug. 2013, doi: 10.1177/0022034513495240.

[29] A. Kassem, C. Lindholm, and U. H. Lerner, “Toll-Like receptor 2 stimulation of osteoblasts mediates staphylococcus aureus induced bone resorption and osteoclastogenesis through enhanced RANKL,” PLoS One, vol. 11, no. 6, Jun. 2016, doi: 10.1371/journal.pone.0156708.

[30] A. N. Ben-awadh et al., “Parathyroid hormone receptor signaling induces bone resorption in the adult skeleton by directly regulating the RANKL gene in osteocytes,” Endocrinology, vol. 155, no. 8, pp. 2797–2809, 2014, doi: 10.1210/en.2014-1046.

[31] P. Curtin, H. Youm, and E. Salih, “Three-dimensional cancer-bone metastasis model using ex-vivo co-cultures of live calvarial bones and cancer cells,” Biomaterials, vol. 33, no. 4, pp. 1065– 1078, Feb. 2012, doi: 10.1016/j.biomaterials.2011.10.046.

[32] M. L. de Vries-van Melle, E. W. Mandl, N. Kops, W. J. L. M. Koevoet, J. A. N. Verhaar, and G. J. V. M. van Osch, “An Osteochondral Culture Model to Study Mechanisms Involved in Articular Cartilage Repair,” Tissue Eng Part C Methods, vol. 18, no. 1, pp. 45–53, Jan. 2012, doi: 10.1089/ten.tec.2011.0339.

[33] S. Ansari, K. Ito, and S. Hofmann, “Cell Sources for Human In vitro Bone Models,” Curr Osteoporos Rep, vol. 19, no. 1, pp. 88–100, Feb. 2021, doi: 10.1007/S11914-020-00648-6/TABLES/2.

[34] B. Kirstein, U. Grabowska, B. Samuelsson, M. Shiroo, T. J. Chambers, and K. Fuller, “A novel assay for analysis of the regulation of the function of human osteoclasts,” J Transl Med, vol. 4, no. 1, pp. 1–9, Nov. 2006, doi: 10.1186/1479-5876-4-45/FIGURES/5.

[35] M. Sørensen et al., “Characterization of osteoclasts derived from CD14+ monocytes isolated from peripheral blood,” J Bone Miner Metab, vol. 25, pp. 36–45, 2007, doi: 10.1007/s00774-006-0725-9.

[36] V. Shalhoub et al., “Characterization of osteoclast precursors in human blood,” Br J Haematol, vol. 111, no. 2, pp. 501–512, Nov. 2000, doi: 10.1111/J.1365-2141.2000.02379.X.

[37] M. Susa, N. H. Luong-Nguyen, D. Cappellen, N. Zamurovic, and R. Gamse, “Human primary osteoclasts: in vitro generation and applications as pharmacological and clinical assay,” J Transl Med, vol. 2, p. 6, Mar. 2004, doi: 10.1186/1479-5876-2-6.

[38] H. K. Yong, J. H. Jun, M. W. Kyung, H. M. Ryoo, G. S. Kim, and J. H. Baek, “Dexamethasone inhibits the formation of multinucleated osteoclasts via down-regulation of beta3 integrin expression,” Arch Pharm Res, vol. 29, no. 8, pp. 691–698, Aug. 2006, doi: 10.1007/BF02968254.

[39] J. Li et al., “Different bone remodeling levels of trabecular and cortical bone in response to changes in Wnt/β-catenin signaling in mice,” Journal of Orthopaedic Research, vol. 35, no. 4, pp. 812–819, Apr. 2017, doi: 10.1002/JOR.23339.

[40] Y. C. Teh, J. L. Ding, L. G. Ng, and S. Z. Chong, “Capturing the fantastic voyage of monocytes through time and space,” Front Immunol, vol. 10, no. MAR, p. 430534, Apr. 2019, doi: 10.3389/FIMMU.2019.00834/BIBTEX.

[41] S. C. Manolagas, “Birth and Death of Bone Cells: Basic Regulatory Mechanisms and Implications for the Pathogenesis and Treatment of Osteoporosis,” Endocr Rev, vol. 21, no. 2, pp. 115–137, Apr. 2000, doi: 10.1210/EDRV.21.2.0395.

[42] M. Kotani et al., “Systemic Circulation and Bone Recruitment of Osteoclast Precursors Tracked by Using Fluorescent Imaging Techniques,” The Journal of Immunology, vol. 190, no. 2, pp. 605–612, Jan. 2013, doi: 10.4049/JIMMUNOL.1201345.

[43] Y. Zhou, H. W. Deng, and H. Shen, “Circulating monocytes: an appropriate model for bone-related study,” Osteoporosis International 2015 26:11, vol. 26, no. 11, pp. 2561–2572, Jul. 2015, doi: 10.1007/S00198-015-3250-7.

[44] N. Udagawa et al., “Origin of osteoclasts: mature monocytes and macrophages are capable of differentiating into osteoclasts under a suitable microenvironment prepared by bone marrow-derived stromal cells.,” Proc Natl Acad Sci U S A, vol. 87, no. 18, p. 7260, 1990, doi: 10.1073/PNAS.87.18.7260.

[45] N. Kurihara, C. Chenu, M. Miller, C. Civin, and G. D. Roodman, “Identification of committed mononuclear precursors for osteoclast-like cells formed in long term human marrow cultures,” Endocrinology, vol. 126, no. 5, pp. 2733–2741, 1990, doi: 10.1210/ENDO-126-5-2733.

[46] A. Schwab et al., “Ex vivo culture platform for assessment of cartilage repair treatment strategies,” ALTEX, vol. 34, no. 2, pp. 267–277, May 2017, doi: 10.14573/altex.1607111.

[47] T. Klüter et al., “An *Ex Vivo* Bone Defect Model to Evaluate Bone Substitutes and Associated Bone Regeneration Processes,” Tissue Eng Part C Methods, vol. 26, no. 1, pp. 56–65, Jan. 2020, doi: 10.1089/ten.tec.2019.0274.

[48] P. Christen, S. Boutroy, R. Ellouz, R. Chapurlat, and B. van Rietbergen, “Least-detectable and age-related local in vivo bone remodelling assessed by time-lapse HR-pQCT,” PLoS One, vol. 13, no. 1, p. e0191369, Jan. 2018, doi: 10.1371/journal.pone.0191369.

[49] P. Christen et al., “Bone remodelling in humans is load-driven but not lazy,” Nature Communications 2014 5:1, vol. 5, no. 1, pp. 1–5, Sep. 2014, doi: 10.1038/ncomms5855.

[50] R. L. Jilka, “The Relevance of Mouse Models for Investigating Age-Related Bone Loss in Humans,” J Gerontol A Biol Sci Med Sci, vol. 68, no. 10, p. 1209, Oct. 2013, doi: 10.1093/GERONA/GLT046.

[51] D. Zhao et al., “Connexin 43 Channels in Osteocytes Regulate Bone Responses to Mechanical Unloading,” Front Physiol, vol. 11, p. 299, Mar. 2020, doi: 10.3389/FPHYS.2020.00299/BIBTEX.

[52] P. Zhang, K. Hamamura, and H. Yokota, “A Brief Review of Bone Adaptation to Unloading,” Genomics Proteomics Bioinformatics, vol. 6, no. 1, pp. 4–7, Jan. 2008, doi: 10.1016/S1672-0229(08)60016-9.

[53] H. Kondo et al., “Unloading induces osteoblastic cell suppression and osteoclastic cell activation to lead to bone loss via sympathetic nervous system,” Journal of Biological Chemistry, vol. 280, no. 34, pp. 30192–30200, Aug. 2005, doi: 10.1074/jbc.M504179200.

[54] F. Cosmi, N. Steimberg, and G. Mazzoleni, “A mesoscale study of the degradation of bone structural properties in modeled microgravity conditions,” J Mech Behav Biomed Mater, vol. 44, pp. 61–70, Apr. 2015, doi: 10.1016/j.jmbbm.2015.01.002.

[55] S. R. Tuljapurkar et al., “Changes in human bone marrow fat content associated with changes in hematopoietic stem cell numbers and cytokine levels with aging,” J Anat, vol. 219, no. 5, p. 574, Nov. 2011, doi: 10.1111/J.1469-7580.2011.01423.X.

[56] O. Kandarakov, A. Belyavsky, and E. Semenova, “Bone Marrow Niches of Hematopoietic Stem and Progenitor Cells,” Int J Mol Sci, vol. 23, no. 8, Apr. 2022, doi: 10.3390/IJMS23084462.

[57] T. H. Ambrosi et al., “Adipocyte Accumulation in the Bone Marrow during Obesity and Aging Impairs Stem Cell-Based Hematopoietic and Bone Regeneration,” Cell Stem Cell, vol. 20, no. 6, pp. 771–784.e6, Jun. 2017, doi: 10.1016/J.STEM.2017.02.009.

[58] D. R. Halloran et al., “Differentiation of Cells Isolated from Human Femoral Heads into Functional Osteoclasts,” J Dev Biol, vol. 10, no. 1, Mar. 2022, doi: 10.3390/JDB10010006.

[59] E. Kylmäoja et al., “Peripheral blood monocytes show increased osteoclast differentiation potential compared to bone marrow monocytes,” Heliyon, vol. 4, no. 9, p. e00780, Sep. 2018, doi: 10.1016/J.HELIYON.2018.E00780.

[60] T. Suda, N. Takahashi, N. Udagawa, E. Jimi, M. T. Gillespie, and T. J. Martin, “Modulation of Osteoclast Differentiation and Function by the New Members of the Tumor Necrosis Factor Receptor and Ligand Families,” Endocr Rev, vol. 20, no. 3, pp. 345–357, Jun. 1999, doi: 10.1210/EDRV.20.3.0367.

[61] A. Papadimitropoulos et al., “A 3D in vitro bone organ model using human progenitor cells,” Eur Cell Mater, vol. 21, pp. 445–458, 2011, doi: 10.22203/ECM.V021A33.

[62] J. J. Cody, A. A. Rivera, J. Liu, J. M. Liu, J. T. Douglas, and X. Feng, “A simplified method for the generation of human osteoclasts in vitro,” Int J Biochem Mol Biol, vol. 2, no. 2, p. 183, 2011, Accessed: Dec. 12, 2022. [Online]. Available: /pmc/articles/PMC3180092/

[63] A. Bernhardt, M. Schumacher, and M. Gelinsky, “Formation of osteoclasts on calcium phosphate bone cements and polystyrene depends on monocyte isolation conditions,” Tissue Eng Part C Methods, vol. 21, no. 2, pp. 160–170, Feb. 2015, doi: 10.1089/TEN.TEC.2014.0187/ASSET/IMAGES/LARGE/FIGURE9.JPEG.

[64] D. Steller, A. Scheibert, T. Sturmheit, and S. G. Hakim, “Establishment and validation of an in vitro co-culture model for oral cell lines using human PBMC-derived osteoclasts, osteoblasts, fibroblasts and keratinocytes,” Scientific Reports 2020 10:1, vol. 10, no. 1, pp. 1–10, Oct. 2020, doi: 10.1038/s41598-020-73941-0.

[65] T. Akchurin, T. Aissiou, N. Kemeny, E. Prosk, N. Nigam, and S. v. Komarova, “Complex Dynamics of Osteoclast Formation and Death in Long-Term Cultures,” PLoS One, vol. 3, no. 5, p. e2104, May 2008, doi: 10.1371/JOURNAL.PONE.0002104.

[66] D. Abdallah et al., “An optimized method to generate human active osteoclasts from peripheral blood monocytes,” Front Immunol, vol. 9, no. APR, p. 632, Apr. 2018, doi: 10.3389/FIMMU.2018.00632/BIBTEX.

[67] B. W. M. De Wildt et al., “The Impact of Culture Variables on a 3D Human In Vitro Bone Remodeling Model: A Design of Experiments Approach,” Adv Healthc Mater, p. 2301205, 2023, doi: 10.1002/ADHM.202301205.

[68] A. M. J. Møller et al., “Fusion Potential of Human Osteoclasts In Vitro Reflects Age, Menopause, and In Vivo Bone Resorption Levels of Their Donors—A Possible Involvement of DC-STAMP,” International Journal of Molecular Sciences 2020, Vol. 21, Page 6368, vol. 21, no. 17, p. 6368, Sep. 2020, doi: 10.3390/IJMS21176368.

[69] D. M. H. Merrild et al., “Pit- and trench-forming osteoclasts: a distinction that matters,” Bone Research 2015 3:1, vol. 3, no. 1, pp. 1–11, Dec. 2015, doi: 10.1038/boneres.2015.32.

[70] J. M. W. Quinn, S. Neale, Y. Fujikawa, J. D. McGee, and N. A. Athanasou, “Human osteoclast formation from blood monocytes, peritoneal macrophages, and bone marrow cells,” Calcif Tissue Int, vol. 62, no. 6, pp. 527–531, 1998, doi: 10.1007/S002239900473.

[71] T. L. Andersen et al., “A Physical Mechanism for Coupling Bone Resorption and Formation in Adult Human Bone,” Am J Pathol, vol. 174, no. 1, p. 239, 2009, doi: 10.2353/AJPATH.2009.080627.

[72] A. Muto et al., “Lineage-committed osteoclast precursors circulate in blood and settle down into bone,” J Bone Miner Res, vol. 26, no. 12, pp. 2978–2990, Dec. 2011, doi: 10.1002/JBMR.490.

[73] F. Xu and S. L. Teitelbaum, “Osteoclasts: New Insights,” Bone Research 2013 1:1, vol. 1, no. 1, pp. 11–26, Mar. 2013, doi: 10.4248/br201301003.

[74] C. S. Soltanoff, S. Yang, W. Chen, and Y. P. Li, “Signaling Networks that Control the Lineage Commitment and Differentiation of Bone Cells,” Crit Rev Eukaryot Gene Expr, vol. 19, no. 1, p. 1, 2009, doi: 10.1615/CRITREVEUKARGENEEXPR.V19.I1.10.

[75] A. Bernhardt, K. Koperski, M. Schumacher, and M. Gelinsky, “Relevance of osteoclast-specific enzyme activities in cell-based in vitro resorption assays,” Eur Cell Mater, vol. 33, pp. 28–42, Jan. 2017, doi: 10.22203/ECM.V033A03.

[76] M. J. Stoddart, P. I. Furlong, A. Simpson, C. M. Davies, and R. G. Richards, “A comparison of non-radioactive methods for assessing viability in ex vivo cultured cancellous bone: Technical note,” Eur Cell Mater, vol. 12, pp. 16–25, 2006, doi: 10.22203/eCM.v012a02.

[77] S. Zankovic et al., “A method for the evaluation of early osseointegration of implant materials ex vivo: Human bone organ model,” Materials, vol. 14, no. 11, p. undefined-undefined, Jun. 2021, doi: 10.3390/MA14113001.

[78] G. Gu, M. Mulari, Z. Peng, T. A. Hentunen, and H. K. Väänänen, “Death of osteocytes turns off the inhibition of osteoclasts and triggers local bone resorption,” Biochem Biophys Res Commun, vol. 335, no. 4, pp. 1095–1101, Oct. 2005, doi: 10.1016/j.bbrc.2005.06.211.

[79] S. J. Morrison and D. T. Scadden, “The bone marrow niche for haematopoietic stem cells,” Nature, vol. 505, no. 7483, p. 327, Jan. 2014, doi: 10.1038/NATURE12984.

[80] N. Baryawno et al., “A cellular taxonomy of the bone marrow stroma in homeostasis and leukemia,” Cell, vol. 177, no. 7, p. 1915, Jun. 2019, doi: 10.1016/J.CELL.2019.04.040.

[81] N. Ahamad, P. C. Rath, N. Ahamad, and P. C. Rath, “Bone Marrow Stem Cells, Aging, and Age-Related Diseases,” *Models*, Molecules and Mechanisms in Biogerontology, pp. 321–352, 2019, doi: 10.1007/978-981-13-3585-3_15.

[82] C. M. Hoffman, J. Han, and L. M. Calvi, “Impact of aging on bone, marrow and their interactions,” Bone, vol. 119, pp. 1–7, Feb. 2019, doi: 10.1016/J.BONE.2018.07.012.

[83] M. Stauber and R. Müller, “Age-related changes in trabecular bone microstructures: Global and local morphometry,” Osteoporosis International, vol. 17, no. 4, pp. 616–626, Apr. 2006, doi: 10.1007/S00198-005-0025-6/FIGURES/4.

[84] C. Wang et al., “Bone Microstructure and Regional Distribution of Osteoblast and Osteoclast Activity in the Osteonecrotic Femoral Head,” PLoS One, vol. 9, no. 5, p. e96361, May 2014, doi: 10.1371/JOURNAL.PONE.0096361.

[85] A. M. J. Møller et al., “Aging and menopause reprogram osteoclast precursors for aggressive bone resorption,” Bone Research 2020 8:1, vol. 8, no. 1, pp. 1–11, Jul. 2020, doi: 10.1038/s41413-020-0102-7.

[86] D. v. Novack, “Estrogen and Bone: Osteoclasts Take Center Stage,” Cell Metab, vol. 6, no. 4, pp. 254–256, Oct. 2007, doi: 10.1016/j.cmet.2007.09.007.

[87] R. K. Fuchs, M. E. Kersh, J. Carballido-Gamio, W. R. Thompson, J. H. Keyak, and S. J. Warden, “Physical activity for strengthening fracture prone regions of the proximal femur,” Curr Osteoporos Rep, vol. 15, no. 1, p. 43, Feb. 2017, doi: 10.1007/S11914-017-0343-6.

[88] B. Li and R. M. Aspden, “Composition and Mechanical Properties of Cancellous Bone from the Femoral Head of Patients with Osteoporosis or Osteoarthritis,” Journal of Bone and Mineral Research, vol. 12, no. 4, pp. 641–651, Apr. 1997, doi: 10.1359/JBMR.1997.12.4.641.

[89] M. W. A. Kleuskens, C. C. van Donkelaar, L. M. Kock, R. P. A. Janssen, and K. Ito, “An ex vivo human osteochondral culture model,” Journal of Orthopaedic Research, vol. 39, no. 4, p. 871, Apr. 2021, doi: 10.1002/JOR.24789.

[90] X. Zhu, Y. T. Chan, P. S. H. Yung, R. S. Tuan, and Y. Jiang, “Subchondral Bone Remodeling: A Therapeutic Target for Osteoarthritis,” Front Cell Dev Biol, vol. 8, p. 1887, Jan. 2021, doi: 10.3389/FCELL.2020.607764/BIBTEX.

