## Supplementary information for "Integration of Osteoclastogenesis through addition of PBMCs in Human Osteochondral Explants cultured *Ex vivo*"

### Donor variability observed between explants

Cross-sections of osteochondral explants stained for H&E revealed differences between explant donors regarding cell abundancy in the bone marrow (**Figure S1, S2A-E**). Donor variability was already apparent at the start of culture where donor 2 demonstrated active bone remodeling throughout the explant (**Figure S1A-E**) and donor 4 appeared to be less active in bone remodeling consisting mainly of fat tissue (**Figure S1F-J**). At the end of culture, marrow contained mainly fat tissue for donor 4 (**Figure 5**), whereas donor 2 demonstrated the presence of other cell types in addition to adipocytes observed from the clusters of intact cell nuclei that were found in the marrow (**Figure S2B-E**). The other 2 donors, 1 and 3, contained marrow cellularity and fat tissue in amounts ranging between donor 2 and 4 (data not shown).

To have a negative control in which cell activity was missing, explants were devitalized by 3 freeze-thaw cycles to induce cell death. Cross-sections of devitalized explants stained with H&E confirmed absence of intact nuclei in the trabecular spaces and absence of osteocytes in the lacunae (**Figure S2F-H**).

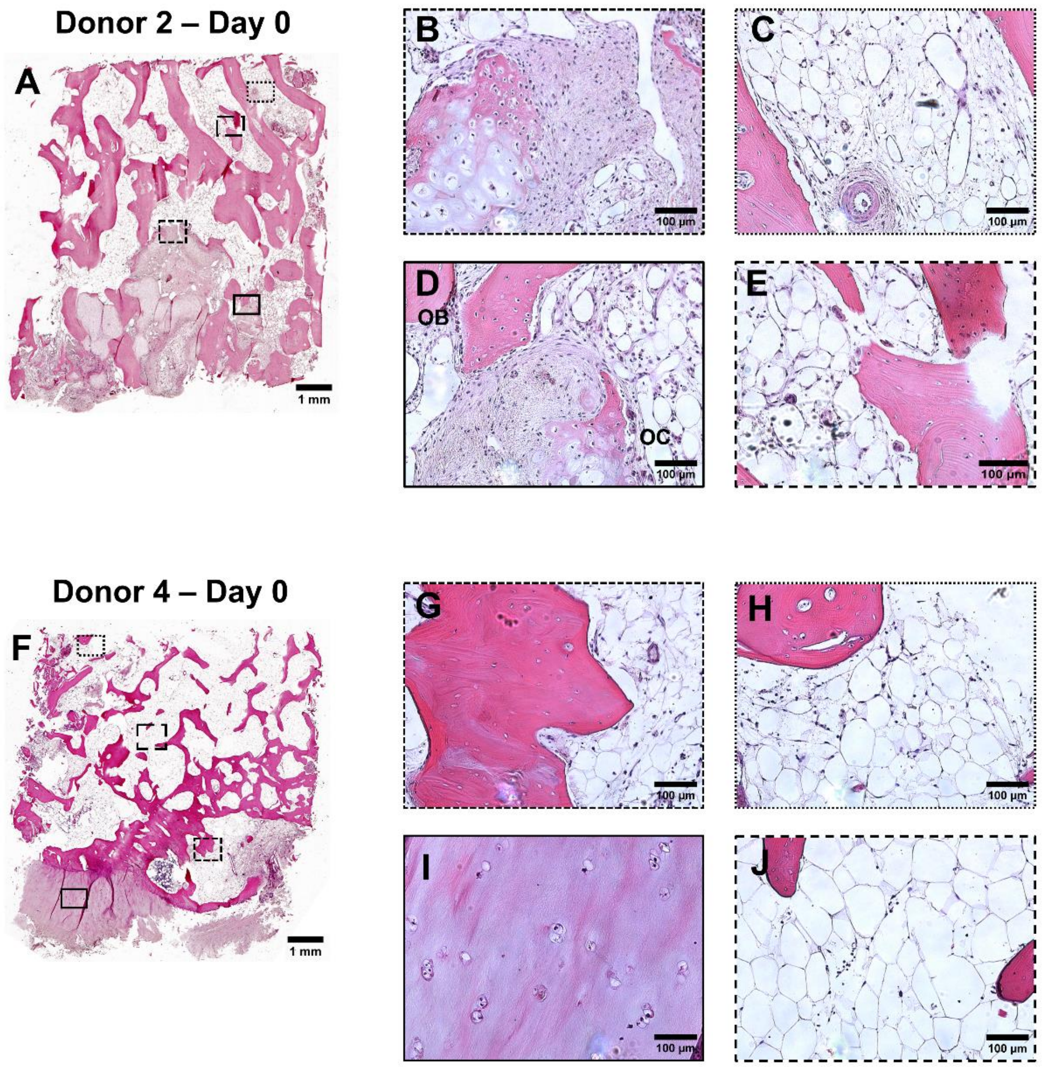


**Figure S1** – Active bone remodeling was observed in osteochondral explants directly after isolation (day 0) stained with H&E (A). Matrix deposition and embedding by osteoblasts (B and D) and resorption by osteoclasts (D-E) was revealed. A different donor demonstrated variability between donors from the start of culture (F-J). No active bone remodeling was seen (G-H) on the surface of trabeculae. Chondrocytes were visualized in cartilage (I) and bone marrow consisted predominantly of adipocytes (J). Abbreviations: osteoblasts (OB) and osteoclasts (OC).

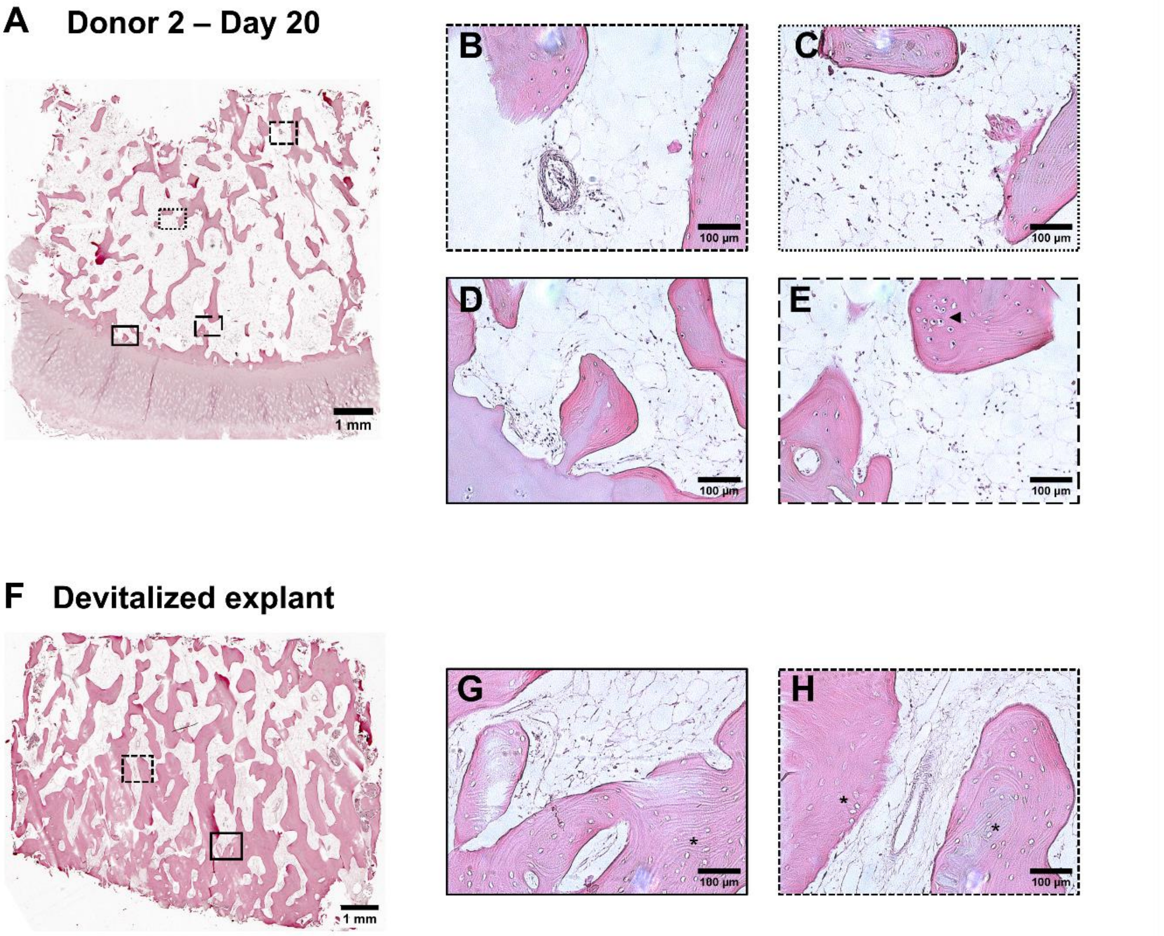


**Figure S2** - Variability between explants donors was observed regarding bone marrow cells. Overview of cross section of osteochondral explants at the end of culture, day 20, stained with H&E (A). High cellular abundance in marrow cavities throughout entire bone part (B-E). As a negative control, explants were devitalized with multiple freeze-thaw cycles before 20-day culture (F). Cellular structures with clear nuclei were absent in bone marrow (G) and osteocytes in lacunae were missing (indicated with *) (H).

### Osteoclasts were present in explants at the start of culture

In addition to immunofluorescent staining for CD61, sections of osteochondral explants were stained for osteoclasts marker TRAP. Staining was performed on samples that were fixed directly after isolation and samples that were cultured for 20 days. Directly after isolation, osteoclasts were found as large multinucleated cells, positive for CD61 and TRAP, forming a border on the bone surface (**Figure S3A-B)**. At the end of culture, TRAP positive multinucleated cells were observed upon addition of PBMCs (**Figure S3C, z-stack movie 3D**). These osteoclast-like cells were not located in clusters but appeared randomly as individual cells throughout the explant.

### Immunofluorescent visualization of osteoclasts with TRAP

Prepared paraffin sections were deparaffinized, hydrated and stained for DAPI and TRAP a marker present in osteoclasts. Briefly, sections were immersed in 1x citrate buffer (S1699, Dako) overnight at 60°C for antigen retrieval. After a washing step in 0.05% Tween (Tween 20, 822184, Merck) in PBS, samples were blocked in 5% normal goat serum in PBS for 1 hour. Primary antibody solution (orb248939, 1:100) was incubated overnight at 4°C. Secondary antibody solution (A21240, 1:200) was incubated for 1 hour at RT followed by DAPI staining, 0.1 µg/ml for 10 min. Images were acquired with a laser scanning microscope (Leica TCS SP8X).

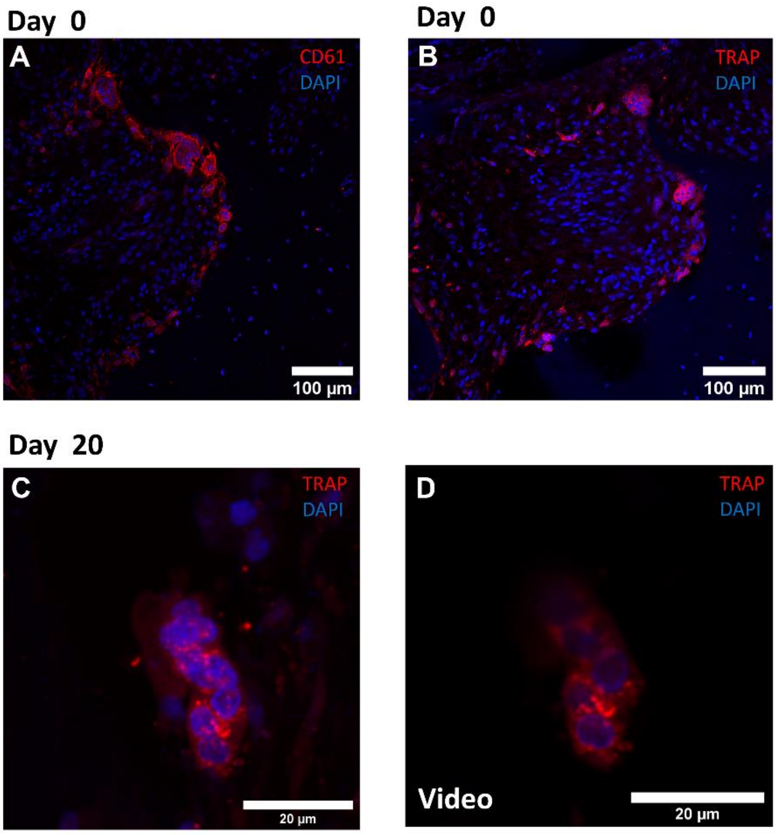


**Figure S3** - Explants revealed multinucleated (dapi, blue) osteoclasts-like cells, observed from osteoclast markers CD61 (A, red) and TRAP (B, red), directly after isolation, day 0. Projection of z-stack of osteoclast on day 20 stained for TRAP (red) and nuclei (blue) (C). Movie of confocal z-stack of osteoclast on day 20 stained for TRAP (red) and nuclei (blue) (D). Abbreviations: peripheral blood mononuclear cells (PBMCs), tartrate resistant acid phosphatase (TRAP).

### Considerations with interpretation of µCT outcomes

Registration of µCT images was used to analyze bone remodeling. Although its power as a tool to detect and quantify areas of bone gain and loss, caution was needed for evaluation and interpretation of gain and loss numbers. The combination of quantitative numbers with visualization of bone gain and loss revealed phenomena that need to be considered for interpretation of remodeling data. A first point of consideration was movement of bone parts. The visualization of registered µCT images showed that small pieces of bone debris, predominantly on the outer surface of the explant, were moving during culture and therefore appeared in both channels of bone gain and loss (**Figure S4A-B**). Furthermore, devitalized explants that lack cell activity, showed to gain mineralized volume at the outer surface (**Figure S4E**).

Morphometric parameters were determined after segmentation at a threshold of 450 mgHA/cm$^3$ using a Gaussian filter with filter support 1 and sigma 0.8. BV/TV and BS/TV were determined from µCT software. The gain in mineralized volume was also represented in the numbers of bone volume/total volume (BV/TV) and bone surface/total surface (BS/TV) where all explants showed an increase in bone volume during the culture period (**Figure S4C-D**). This suggested that there was a gain in mineralized volume independent of cellular activity, possibly (partly) caused by precipitation of calcium and phosphate present in the culture medium. Moreover, the devitalized explants were also used to evaluate noise, which is important to consider in evaluation of remodeling processes. Noise was detected as speckles of gain and loss throughout the devitalized explants (**Figure S4F-G**). Considering the presence of gain and loss speckles located inside trabeculae where no bone cell activity was expected, it was likely that this can be attributed to noise. Also, the presence of both gain and loss speckles in comparable amounts, as observed by eye, supported the hypothesis that this was noise.

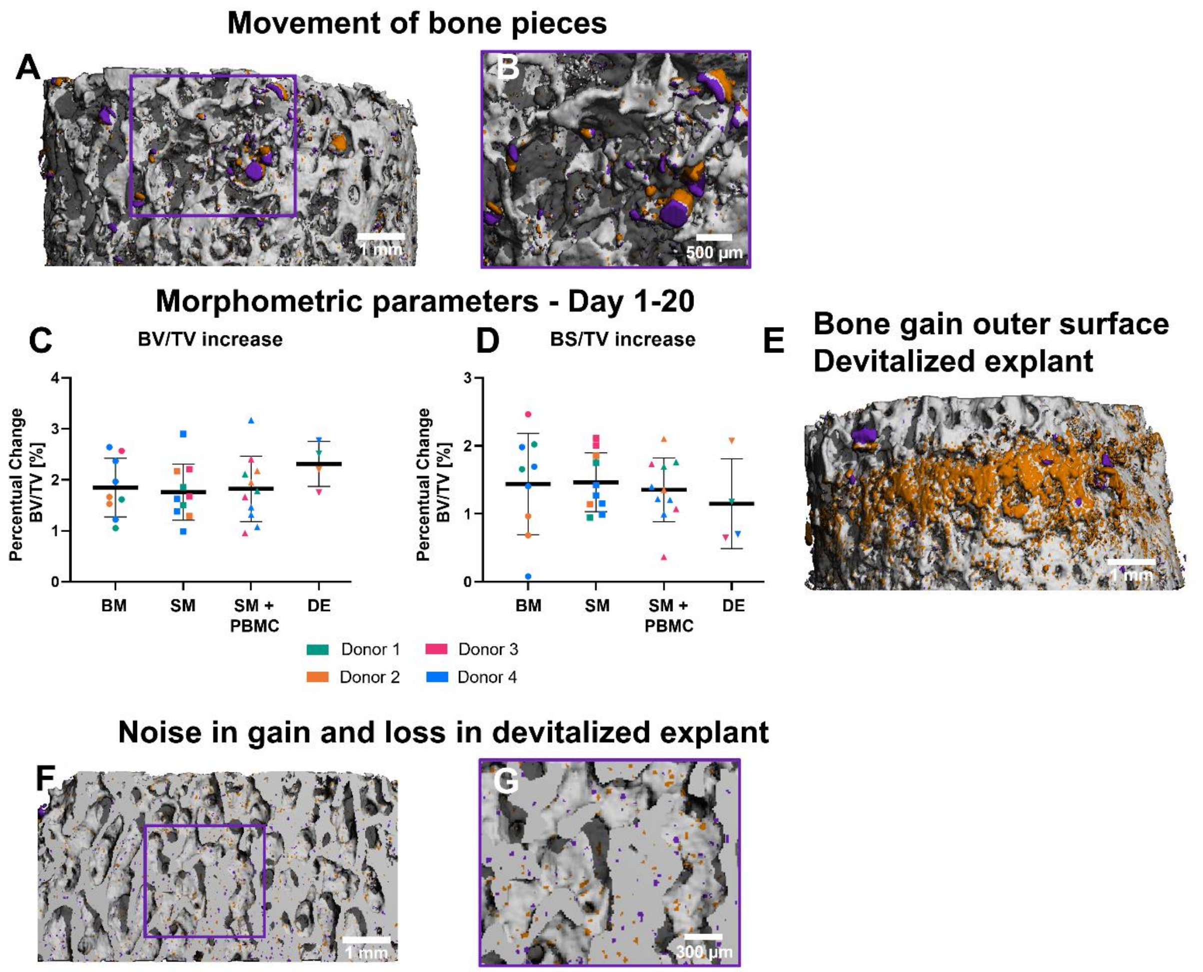


**Figure S4** – Movement of pieces of bone at the outer surface of explants appeared in bone gain (orange) and loss (purple) (A-B). BV/TV (C) and BS/TV (D) increased in all explants over the culture period. Mineralized volume gained at the outer surface of a devitalized explant (E). Devitalized explant, which lacks cell activity, demonstrated gain (orange) and loss (loss) within trabeculae (F-G). Data represents mean ± SD. Abbreviations: peripheral blood mononuclear cells (PBMCs), base medium (BS), supplemented medium (SM), supplemented medium + PBMCs (SM + PBMC), devitalized explant (DE).
